# NeuroMark dFNC: Transferable Dynamic-State Templates for Subject-Specific and Frequency-Resolved Chronnectomics

**DOI:** 10.64898/2026.10.12.758630

**Authors:** M. Moein Esfahani, Masoud Seraji, Sepehr Salem, Kyle M. Jensen, Zening Fu, Tulay Adalı, Yu-Ping Wang, Vince Calhoun

## Abstract

Dynamic functional network connectivity (dFNC) commonly identifies recurring connectivity states from resting-state fMRI, but states derived independently in each dataset lack guaranteed correspondence across studies and cannot be directly estimated in new participants. We introduce NeuroMark dFNC, which extends the NeuroMark principle of prior-guided network estimation to dynamic connectivity by using reproducible reference states as priors for estimating corresponding subject-specific states. Reference-state templates were derived from UK Biobank using windowed functional network connectivity and time-frequency moments (TFMs), with split-sample reproducibility evaluated for k-means, independent component analysis (ICA), and principal component analysis (PCA) template-generation approaches. Constrained ICA then used the templates to estimate subject-specific state patterns and continuous loadings in independent FBIRN schizophrenia and ADNI Alzheimer’s disease cohorts. ICA loadings were calibrated by their contribution to reconstruction of the original feature matrix. Fractional occupancy and conditional mean dwell time were calculated from the state with the largest absolute calibrated loading at each observation. The predefined templates avoided the need for dataset-specific state discovery and post-hoc state matching. In both clinical cohorts, the analysis identified state-dependent, distributed edgewise differences and differences in selected temporal summaries. Nominal frequency-resolved findings varied across states and moment-frequency bins and were treated as exploratory. NeuroMark dFNC provides a common state space for cross-cohort comparison while retaining subject-specific connectivity patterns and continuous state expression.

## 1 Introduction

Resting-state functional magnetic resonance imaging (rs-fMRI) measures spontaneous blood-oxygenation-level-dependent (BOLD) fluctuations and is widely used to study intrinsic functional organization [1]. Functional connectivity summarizes statistical dependence between signals from spatially distinct regions or networks. These estimates vary across individuals and within an individual over time, reflecting a mixture of neural dynamics, physiological effects, motion, and measurement noise. Methods intended for cross-cohort or clinical use must therefore preserve meaningful individual variation while producing comparable features across participants and studies.

Group ICA provides a data-driven approach for identifying common functional brain networks from rs-fMRI [2, 3, 4], but maintaining correspondence of individual network estimates across subjects and datasets can be challenging. Constrained ICA addresses this problem by using spatial priors to guide subject-level decomposition. NeuroMark applies reproducible spatial network templates within this framework, enabling corresponding subject-specific networks to be estimated across datasets while preserving individual variability [5, 6, 7, 8, 9]. However, this standardization has largely been restricted to spatial network estimation, while dynamic connectivity states are still typically defined independently within each dataset.

Beyond static representations of connectivity, the field has increasingly recognized that functional interactions between brain regions are not fixed but are inherently dynamic, fluctuating over various timescales [10, 11],[12]. Dynamic functional network connectivity (dFNC) analysis seeks to characterize these time-varying patterns of inter-network communication, providing a representation of connectivity that captures temporal variation not represented by static connectivity measures. A common dFNC pipeline involves identifying RSNs, calculating time-varying connectivity between them (often using sliding windows), and then employing techniques like clustering or further decomposition to identify recurring transient “brain states” [13].

Most Dynamic functional network connectivity (dFNC) or dynamic functional connectivity (dFC) pipelines define states independently within each dataset. Consequently, a state identified in one cohort is not guaranteed to correspond to a state identified in another, and assigning new participants to an established state space generally requires post-hoc matching or classification. This limits direct replication, cross-cohort comparison, and development of standardized subject-level dynamic features. A transferable framework should therefore define a common state space while allowing each participant to express individualized state patterns and temporal profiles.

Here we extend the NeuroMark framework from spatial networks to dynamic connectivity states. Dynamic-state templates are first derived from an independent reference cohort and then used as priors in constrained ICA to estimate corresponding subject-specific states in new participants. We evaluate this framework using both conventional windowed functional network connectivity (wFNC) and a complementary TFM representation that resolves state expression across time-frequency moments [14]. Reference templates are derived and evaluated using k-means, PCA, and ICA, and all three template sets are provided as reusable priors. We demonstrate template-guided subject-specific estimation using the ICA-derived priors. An overview of template construction and subject-specific state estimation is provided in Figure 1.

**Figure 1:**
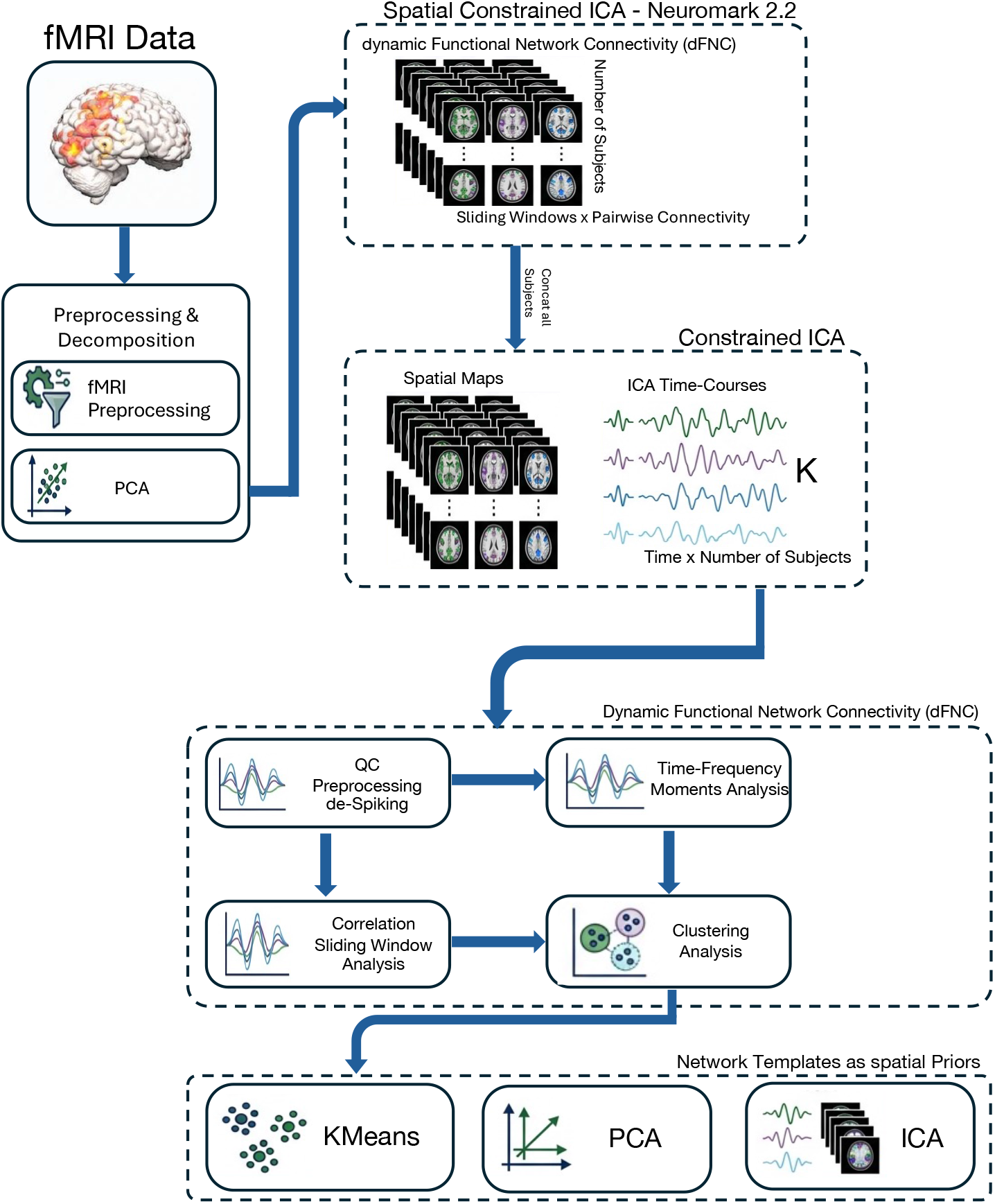
Overview of the NeuroMark dFNC template-construction pipeline. Preprocessed resting-state fMRI data are reduced by PCA and decomposed by spatially constrained ICA using the NeuroMark 2.2 priors to obtain corresponding subject-specific intrinsic connectivity networks and their time courses. After quality control and time-course post-processing, conventional windowed functional network connectivity and frequency-aware time-frequency-moment features are estimated. Features are aggregated across the UK Biobank reference cohort, and k-means, ICA, and PCA are used to derive complementary sets of reproducible dynamic-state priors. All three template sets are provided for reuse; the present study demonstrates constrained subject-specific state estimation using the ICA-derived priors.

**Figure 2:**
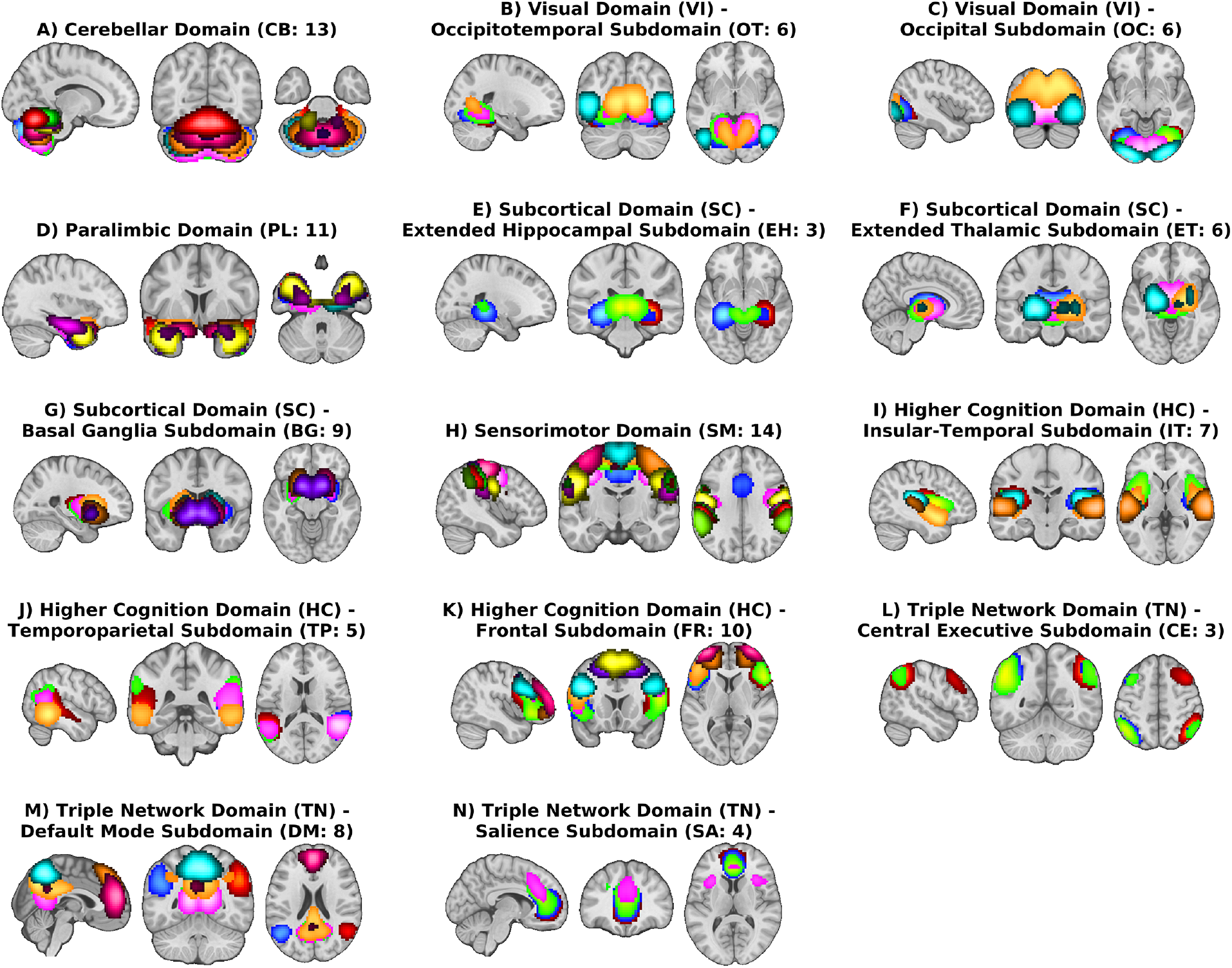
NeuroMark 2.2 multiscale intrinsic connectivity network (ICN) reference template. The template comprises 105 ICNs organized into seven functional domains and 14 subdomains: cerebellar (CB; 13 ICNs), visual– occipitotemporal (VI–OT; 6), visual–occipital (VI–OC; 6), paralimbic (PL; 11), subcortical–extended hippocampal (SC–EH; 3), subcortical–extended thalamic (SC–ET; 6), subcortical–basal ganglia (SC–BG; 9), sensorimotor (SM; 14), higher cognition–insular temporal (HC–IT; 7), higher cognition–temporoparietal (HC–TP; 5), higher cognition–frontal (HC–FR; 10), triple network–central executive (TN–CE; 3), triple network–default mode (TN–DM; 8), and triple network–salience (TN–SA; 4). Colored overlays distinguish the constituent ICNs within each subdomain; the number in parentheses is the number of ICNs represented.

We test three questions: (1) Are the reference-state templates reproducible in independent UK Biobank subsets? (2) Can the same templates be used to estimate corresponding subject-specific states in independent schizophrenia and Alzheimer’s disease cohorts? and (3) Do the resulting state patterns and expression measures retain sensitivity to clinically relevant group differences?

### 1.1 Literature Review

The emergence of time-resolved functional (network) connectivity marked a significant advance from static views, acknowledging the time-varying nature of neural interactions [10, 11, 12]. The predominant method for estimating dFNC has been the sliding-window approach, typically applied to time courses derived from resting-state networks (RSNs) defined by ICA [13, 15]. Within each window, connectivity is estimated, and the resulting series of connectivity matrices are then often subjected to clustering algorithms, most commonly k-means, to identify recurring “brain states”—distinct patterns of whole-brain inter-network communication [13, 16]. Such dFNC states have demonstrated sensitivity to various neurological and psychiatric conditions [16, 17, 18], offering potential for the development of biomarkers. For example, altered dwell times or transitions between specific dFNC states have been reported in schizophrenia [16, 19] and Alzheimer’s disease [20, 21, 22], suggesting that aberrant dynamic functional organization is a key feature of these disorders.

Despite its widespread adoption, the sliding-window k-means (SWK) approach to dFNC has recognized limitations. The choice of window length, step size, and number of clusters (k) can significantly influence the resulting states, potentially affecting replicability [23, 24, 25]. Furthermore, the assumption of stable connectivity within each window may not always be physiologically accurate. These concerns have spurred exploration of alternative dFNC estimation techniques. These include methods that dispense with windowing, such as instantaneous phase-synchrony and dictionary learning approaches [26, 27, 28], co-activation pattern (CAP) analysis, which identifies recurring patterns of regional co-activation [29], and point process analysis [30]. Other innovations include hidden Markov models (HMMs) to infer discrete brain states and their temporal dynamics directly from BOLD data without explicit windowing [31, 32, 33] as well as filterbank approaches to cover a range of windowing/filter choices [34, 35, 28].

While these approaches improve flexibility or temporal resolution, they largely retain dataset-specific state definitions and do not address cross-study comparability or subject-level generalization. At the same time, progress has been made in estimating subject-specific RSNs, which are essential inputs for dFNC. Group ICA can capture population-level RSNs [3], but translating those into accurate individual-level maps with correspondence across studies is not always straightforward. Constrained ICA methods, especially those using spatial priors like the NeuroMark templates [5, 6, 36], have enabled a fully automated approach that can capture individual differences [37, 38]. While such hybrid approaches have made gains in personalizing spatial networks, the next big step is to extend that same level of standardization to the dynamic states themselves.

Time-frequency moments (TFM) provide a complementary dynamic representation by resolving standardized statistical-moment signals jointly over time and moment-space frequency [14]. Both wFNC and TFM were computed from ICN time courses band-pass filtered at 0.01–0.15 Hz. TFM resolves temporal variation in pairwise standardized-moment signals rather than applying additional frequency-band filtering to the ICN time courses. In the present study, pairwise TFMs correspond to the second-order mixed moment **r** = [1, 1].

In addition, a central challenge impeding the clinical translation of dFNC findings is the robust identification and quantification of comparable dFNC states in different individuals and studies. Although group-level analyses can identify common recurring states, it remains difficult to reliably map these group-defined states to new, individual subjects or ensure that states derived from different datasets are truly analogous [39, 40]. Most dFNC studies derive states de novo for each dataset, resulting in state definitions that are not directly comparable across studies and cannot be consistently estimated in new individuals. Without a standardized approach to define and identify states in individuals, the development of generalizable dFNC biomarkers is problematic. There have been limited attempts to address this, for example, by assigning individual dFNC windows to pre-defined group-level state centroids based on similarity [13]. However, such post-hoc assignment may not optimally capture individual deviations or ensure that the assigned state truly reflects the underlying individual dynamics in a constrained and regularized manner. What is largely missing is a framework that explicitly uses well-characterized dFNC state definitions as priors to guide the estimation of these states at the subject level in an automated and consistent fashion.

The development of large-scale neuroimaging resources like the UK Biobank [41, 42], coupled with robust tools for preprocessing and initial network definition like NeuroMark [5, 7], provides an unprecedented opportunity to develop and validate such standardized dFNC approaches. However, leveraging these resources effectively for dFNC requires methods that can overcome the aforementioned limitations of comparability and individual subject generalization.

Spatial priors have enabled corresponding subject-specific networks to be estimated across studies, but dynamic states are still commonly defined separately in each dataset. The present work extends prior-guided estimation to dynamic states and treats them as transferable templates with subject-specific edgewise patterns and expression profiles.

## 2 Materials and Methods

### 2.1 Template Dataset (UK Biobank)

In this study, multiple datasets were used. State templates were derived from the UK Biobank (UKB), a large-scale population-based cohort with standardized imaging protocols and deep phenotyping [43]. Dynamic-state templates were derived from the main 39,342 UK Biobank participants who passed the quality-control criteria described below. UK Biobank was used exclusively as the reference dataset for template estimation and split-sample reproducibility analysis. Participants were randomly divided into two independent subsets of and N=1000 subjects for template replication.

We used a five-state representation for all subsequent analyses. The model order, *K* = 5, was selected using the elbow criterion, supported by prior dFNC literature [16], and fixed before application to FBIRN and ADNI. To assess sensitivity to model order, template reproducibility was also examined across multiple values of *K* and is reported in the Supplementary Materials.

For this work, we used resting-state fMRI (rs-fMRI) data from a subset of UKB participants who passed standard quality control procedures, including checks for excessive motion and imaging artifacts. Imaging data were acquired using harmonized protocols across sites, enabling the derivation of stable and generalizable connectivity patterns. These templates are subsequently used as priors for subject-level estimation in independent datasets, enabling cross-study comparability and individualized analysis within a unified framework.

### 2.2 Application to Two Clinical Datasets

We evaluated the NeuroMark dFNC pipeline in two independent clinical datasets: FBIRN, which included individuals with schizophrenia and healthy controls, and ADNI, which included cognitively normal participants and participants with Alzheimer’s disease.

#### 2.2.1 FBIRN

We next utilized the resulting state templates to analyze the phase III Function Biomedical Informatics Research Network (FBIRN) dataset [44], a multi-site study designed to investigate schizophrenia using harmonized imaging and clinical protocols. After quality control, the FBIRN Phase III sample included 151 participants with schizophrenia and 160 healthy controls acquired across eight sites. The cohort was used as an independent psychiatric test dataset for evaluating transfer of the UKB-derived state templates and testing diagnostic-group differences in subject-specific state measures.

For this study, we used resting-state fMRI data from participants in the FBIRN Phase III cohort, which includes individuals with schizophrenia and matched healthy controls. All data were acquired using standardized 3.0T MRI protocols across sites, reducing inter-scanner variability while preserving realistic multi-site heterogeneity.

The FBIRN dataset enables evaluation of whether state templates derived from UKB can be reliably applied to an independent clinical population. In particular, it allows assessment of (i) the stability of subject-specific state estimation and (ii) the sensitivity of the resulting dynamic connectivity measures to schizophrenia-related differences.

#### 2.2.2 ADNI

We also evaluated generalizability using data from the Alzheimer’s Disease Neuroimaging Initiative (ADNI) [45], a large, multi-site longitudinal study focused on identifying biomarkers of Alzheimer’s disease. ADNI includes participants spanning healthy controls, individuals with mild cognitive impairment (MCI), and individuals with Alzheimer’s disease (AD), providing a spectrum of neurodegenerative progression.

For this work, we used resting-state fMRI data from ADNI participants who passed standard quality control procedures. After quality control, the ADNI sample included 830 cognitively normal participants and 741 participants with Alzheimer’s disease. Imaging data were acquired across multiple sites and scanner platforms (including GE, Siemens, and Philips systems) using standardized acquisition protocols designed to ensure cross-site consistency. ADNI served as an independent neurodegenerative test cohort for evaluating whether the same UKB-derived reference states could be transferred to a population differing substantially in age, disease process, and acquisition characteristics from FBIRN.

The inclusion of ADNI enables evaluation of the proposed framework in a distinct clinical domain with different underlying pathophysiology. This allows us to assess whether the state-template–based approach generalizes beyond psychiatric populations and captures meaningful alterations in dynamic connectivity associated with neurodegenerative processes.

### 2.3 Data Processing

#### 2.3.1 Data Quality Control

Quality control followed the framework of Iraji et al. [8]. Participants were required to have at least 120 retained resting-state fMRI volumes. Mean framewise displacement was required to be below 0.25, and participants were excluded when any volume-to-volume motion transition exceeded 3^*°*^ of rotation or 3 mm of translation in any direction. Registration to the EPI template was assessed for anatomical plausibility. Spatial overlap between each participant’s brain mask and the group mask was required to exceed 80% for the whole brain and separately for the upper and lower 10 image slices. Participants failing any criterion were excluded before ICN estimation.

#### 2.3.2 Data Preprocessing

For the UKB dataset, we obtained preprocessed resting-state functional MRI (rsfMRI) data from the UK Biobank Accelerating Medicines Partnership (AMS) server (https://ams.ukbiobank.ac.uk/ams/). Additional preprocessing procedures were performed to further standardize the imaging data. Specifically, rsfMRI volumes were spatially normalized to the standard echo-planar imaging (EPI) template space, followed by spatial smoothing using a 6-mm full width at half maximum (FWHM) Gaussian kernel to improve the signal-to-noise ratio. To ensure data quality and reliability, subsequent quality-control procedures were applied to the imaging data. The preprocessing procedures are described in detail in the UK Biobank imaging documentation (Section 3.1; https://biobank.ctsu.ox.ac.uk/crystal/crystal/docs/brain_mri.pdf). Briefly, the UKB minimal preprocessing pipeline incorporates several standardized procedures to ensure imaging quality and consistency across participants. Initial preprocessing included motion correction using MCFLIRT, followed by grand-mean intensity normalization of the entire 4D dataset using a single multiplicative scaling factor. Temporal high-pass filtering was subsequently applied using Gaussian-weighted least-squares straight-line fitting (*σ* = 50 s) to reduce low-frequency signal drifts. Geometric distortions associated with echo-planar imaging (EPI) acquisition were corrected through fieldmap-based EPI unwarping and gradient distortion correction (GDC). Structured noise and imaging artifacts were further identified and removed using ICA+FIX denoising, which combines ICA with FMRIB’s ICA-based X-noiseifier (FIX). The FIX classifier was manually trained on 40 UKB rsfMRI datasets and validated using a leave-one-out approach, demonstrating high classification accuracy for both non-artifactual and artifactual components.

The FBIRN and ADNI data were processed with SPM12 [46, 47]. The first 10 volumes were removed, followed by slice-timing correction and rigid-body motion correction. No additional voxelwise nuisance regression was applied before ICA. Images were normalized to Montreal Neurological Institute space using an EPI template, resampled to 3-mm isotropic voxels, and smoothed with a 6-mm FWHM Gaussian kernel, consistent with the NeuroMark processing described by Iraji et al. [8]. Thus, UKB and the two clinical datasets entered the analysis through their respective preprocessing streams but were brought into the same template space before NeuroMark estimation.

### 2.4 Spatially Constrained ICA for Subject-Specific Network Estimation

We applied the NeuroMark framework, a hybrid data-driven approach, to the rsfMRI data to derive subject-specific intrinsic connectivity networks (ICNs) for subsequent functional network connectivity (FNC) analyses. NeuroMark combines a robust predefined spatial network template [48] with spatially constrained independent component analysis (ICA), thereby integrating the advantages of atlas-based and fully data-driven methods. In refrence [48], this template mentioned as neuromark 2.2. Compared with conventional atlas-based approaches, which may be limited by predefined and potentially arbitrary parcellations, the template-guided ICA framework provides greater flexibility in capturing individual functional organization. In contrast to unconstrained data-driven decomposition methods, which can produce components with limited correspondence across subjects or studies, NeuroMark imposes standardized spatial priors to ensure that extracted ICNs remain spatially consistent and directly comparable across datasets. This framework improves the reproducibility and generalizability of large-scale FNC analyses while preserving sensitivity to subject-level variability in network organization, thereby facilitating robust cross-subject and cross-cohort investigations of brain functional connectivity patterns.

The component time courses were detrended to remove linear, quadratic, and cubic trends; the six rigid-body motion parameters and their temporal derivatives were regressed out; outlying time points were identified and removed; and the residual time courses were band-pass filtered at 0.01–0.15 Hz. Windowed FNC was then estimated with a tapered window formed by convolving a 40-TR rectangular window with a Gaussian kernel (*σ* = 3 TRs). Because TR differed across cohorts, the duration of a 40-TR window was dataset specific (32 s in UKB, 80 s in FBIRN, and 120 s in ADNI). A fixed 40-TR window was retained to standardize the nominal number of samples contributing to each connectivity estimate across cohorts, while accepting differences in physical window duration.

### 2.5 Characterizing Dynamic States with Time-Frequency Moments

TFMs estimate standardized statistical moments jointly over time and frequency in moment space [14]. Let *x*_*i*_(*t*) denote the post-processed ICN time courses *i*. A normalized 21-sample Gaussian kernel, truncated at *±* 2.5*σ* with *σ* = 4 samples, was used for both local standardization and subsequent temporal smoothing:

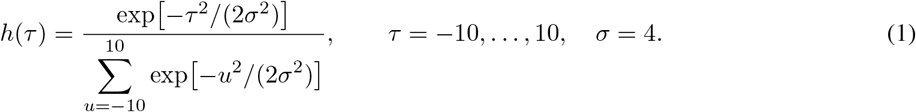

The locally weighted mean, standard deviation, and standardized time course were

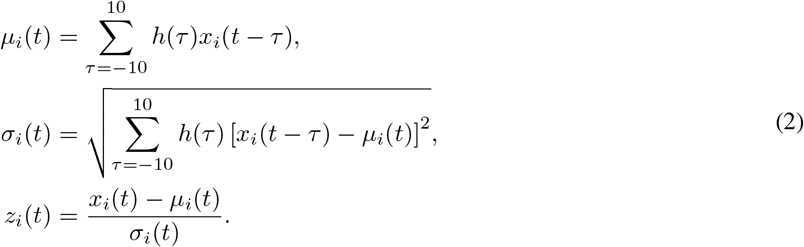

For a moment-order vector **r** = [*r*_1_, …, *r*_*n*_], the instantaneous standardized-moment signal is

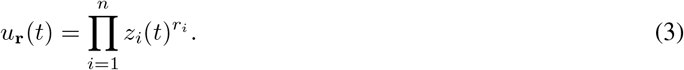

Only pairwise second-order mixed moments were analyzed: **r** = [1, 1] was evaluated for every unordered, off-diagonal ICN pair, with self-moments excluded. For 105 ICNs, this yielded

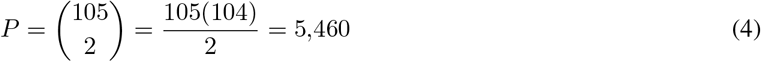

unique pairwise moment signals. Let 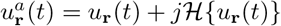 denote the analytic signal, where ℋ{·} is the Hilbert transform. Frequency selection was implemented by single-sideband modulation followed by Gaussian smoothing:

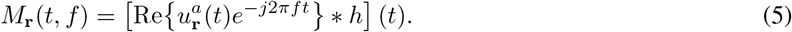

The code-estimated −3 dB cutoff of the Gaussian response defined the nominal bin bandwidth *B*. Because a Gaussian response has nonzero tails, these intervals are nominal −3 dB bands rather than hard filter edges. No separate FIR or IIR band-pass filter was applied; accordingly, no band-pass filter order was specified. No forward–reverse filtering or explicit zero-phase padding was used. The kernel is symmetric and therefore has zero phase relative to its window midpoint. Both Gaussian operations used valid-window processing; no padding was applied, 20 samples were discarded at each end in total, and *T*_out_ = *T*_in_ − 40.

Eight moment-frequency bins were computed in each clinical dataset. For FBIRN (TR= 2 s; Nyquist frequency = 0.25 Hz), the nominal bandwidth was *B* = 0.017 Hz. For ADNI (TR= 3 s; Nyquist frequency *≈* 0.167 Hz), it was *B* = 0.01 Hz. Thus, both datasets used eight bins, but their physical frequency axes differed because of their sampling intervals. These frequencies describe variation in the pairwise standardized-moment signal; they are not frequency bands applied to the original ICN time courses.

The construction and organization of the group-level pairwise TFM feature matrix are summarized in Figure 3.

**Figure 3:**
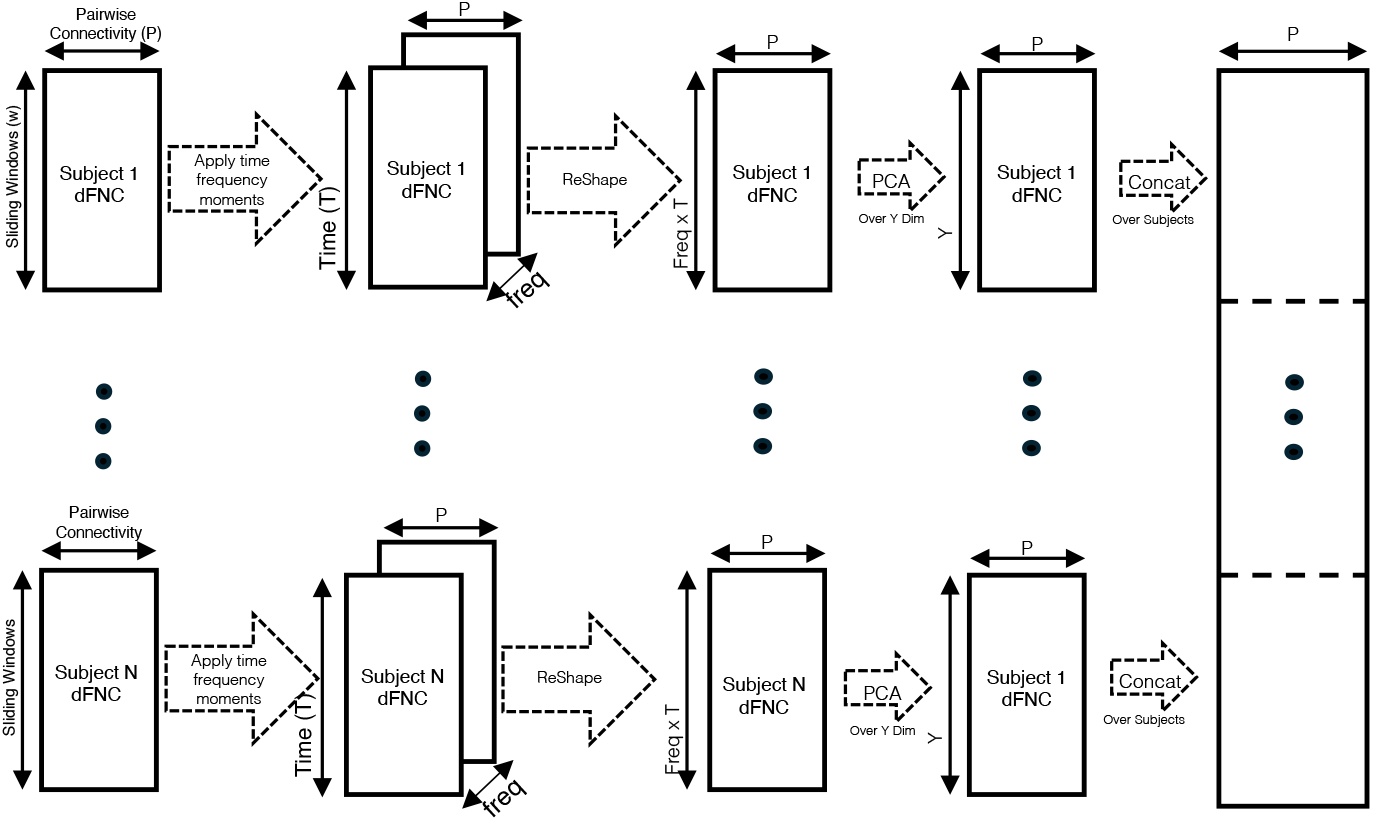
Construction of the group-level pairwise TFM feature matrix. For each subject, locally standardized ICN time courses are multiplied for each of the *P* = 5,460 unique ICN pairs to form the **r** = [1, 1] standardized-moment signals. Single-sideband modulation and Gaussian smoothing resolve these signals over time and eight moment-frequency bins, yielding a time-by-frequency-by-edge representation. The time and frequency dimensions are unfolded, PCA reduces the observation dimension to *Y* retained features while preserving the *P* edges, and the reduced matrices are concatenated across *N* subjects for estimation of the five TFM state templates.

### 2.6 Derivation and Evaluation of Reference Dynamic-State Priors

Reference dynamic-state templates were derived independently for the wFNC and TFM representations using the UK Biobank cohort. For wFNC, each observation was represented by the 5,460 unique pairwise connectivity values among the 105 NeuroMark ICNs. For TFM, the time and moment-frequency dimensions were unfolded to form observations over the same 5,460 ICN-pair feature space; dimensionality reduction was applied along the observation dimension before group-level template estimation. Subject-level feature matrices were then aggregated across the UK Biobank reference cohort.

To evaluate the reproducibility of the reference-state structure, the UK Biobank sample was divided into two independent subsets, denoted Batch A and Batch B. State templates were estimated separately within each subset using three complementary approaches: k-means clustering, independent component analysis (ICA), and principal component analysis (PCA). A five-state model order was used consistently for both the wFNC and TFM representations. As described above, this model order was selected using the elbow criterion and prior dFNC literature [49, 50] and was fixed before evaluation in the clinical cohorts. Corresponding states obtained independently from the two subsets were matched across batches, and reproducibility was quantified using Pearson correlation between the matched edgewise state patterns. This analysis was designed to determine whether similar reference-state structure could be recovered independently of the particular UK Biobank sample used for template estimation.

The three approaches were included both to assess the robustness of the reference-state structure across alternative decomposition methods and to generate complementary sets of reusable dynamic-state priors. ICA-derived templates were selected a priori for the primary subject-specific NeuroMark dFNC analyses. Unlike k-means, which assigns each observation to a single discrete state, ICA provides continuous loadings that permit multiple states to be expressed to different degrees at the same observation. ICA is also directly aligned with the constrained ICA framework used subsequently to estimate corresponding subject-specific state patterns and expression profiles. PCA provides an additional continuous linear representation but does not impose the statistical-independence objective used by the constrained ICA estimation framework.

All three template sets, k-means, PCA, and ICA, are provided as reusable reference priors for future applications and are available through Supplemental Data. In the present study, however, we demonstrate subject-specific template-guided estimation using the ICA-derived priors. ICA was selected for this demonstration because, unlike k-means, it provides continuous state loadings that allow multiple states to be expressed to different degrees at each observation and because its decomposition objective is most directly aligned with the constrained ICA framework used for subject-specific estimation. PCA likewise provides a continuous representation and may be used as a reference prior, but it does not impose the statistical-independence objective used in the present constrained ICA implementation. The choice to demonstrate the ICA priors was methodological and was made independently of group differences in FBIRN or ADNI.

### 2.7 Constrained dFNC Estimation Using cICA

To obtain corresponding subject-specific dynamic states, we extended the multivariate-objective optimization ICA with reference (MOO-ICAR) framework used in NeuroMark spatial ICA to connectivity features [38, 5]. For subject *s* and representation *q* ∈ {wFNC, TFM}, let 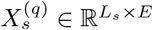 denote the subject-level feature matrix, where *E* is the number of unique ICN-pair edges and *L*_*s*_ indexes temporal windows for wFNC or time–frequency observations for TFM. The corresponding set of *K* group-level state templates is denoted by 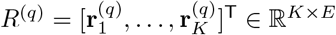.

After centering, dimension reduction, and whitening, 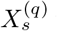 was represented by 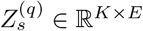. MOO-ICAR estimated the subject-specific state matrix

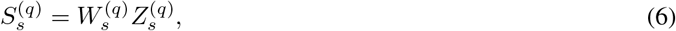

where the *k*th row, 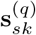, is the subject-specific connectivity pattern corresponding to template 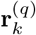. For each state, MOO-ICAR jointly maximized statistical independence and correspondence to its predefined template:

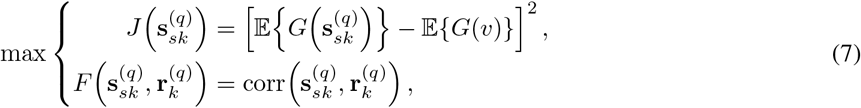

subject to unit-variance and mutual-decorrelation constraints. Here, *G*(*·*) is the nonquadratic function used to approximate negentropy and *v* is a standardized Gaussian variable. Back-reconstruction yielded a loading matrix 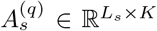 such that 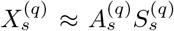. Rows of 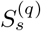 are subject-specific state patterns, and columns of 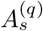 describe their expression across observations. The template-similarity objective establishes state correspondence across subjects and datasets, whereas the independence objective allows the subject-specific patterns to deviate from their group templates.

Because the scale of an ICA component is not identifiable, the raw loadings were calibrated against the original feature matrix before states were ranked. Let **a**_*sk*_ be column *k* of 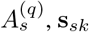 be row *k* of 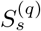, and *C*_*sk*_ = **a**_*sk*_ **s**_*sk*_ be the rank-one contribution of state *k*. We estimated one coefficient per state by

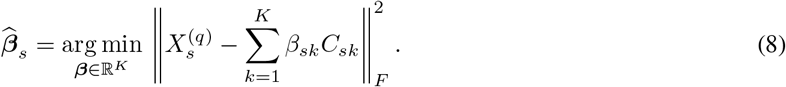

The calculation used the *K × K* Gram matrix and *K*-vector

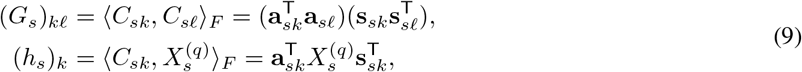

and solved 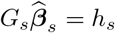 without constructing a vectorized design matrix. This calculation is algebraically equivalent to ordinary least squares on vec 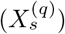; comparison with the explicitly vectorized solution for one participant gave a maximum absolute coefficient difference of 3.35 *×* 10^−14^. The calibrated loading was

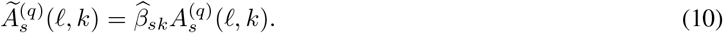

For TFM, *ℓ* indexes the Cartesian product of temporal windows and moment-frequency bins. The calibrated loadings were therefore refolded as 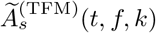. This representation preserves a common edgewise state pattern while allowing its expression to vary over time and moment frequency.

### 2.8 Calibrated State Ranking and Temporal Metrics

Constrained ICA estimates a continuous expression coefficient for each of the *K* states at every observation rather than imposing a hard state label. Discrete state labels were introduced only to calculate fractional occupancy (FO) and conditional mean dwell time (MDT). Because the sign of an ICA component is arbitrary, states were ranked by absolute calibrated loading. For the TFM representation, the state at rank *r* ∈ {1, 2} was

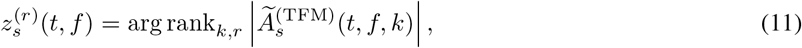

where *r* = 1 and *r* = 2 denote the largest and second-largest contributions, respectively. Ties, which have probability zero for continuous loadings, were resolved by state index. The corresponding wFNC label omits *f*. The primary analyses and Figures 10 and 14 use *r* = 1. Figure 4 illustrates how continuous calibrated state-expression time courses are converted into primary-state assignments and subsequently summarized using FO and conditional MDT. The illustration shows one moment-frequency bin and uses the primary state, *r* = 1.

**Figure 4:**
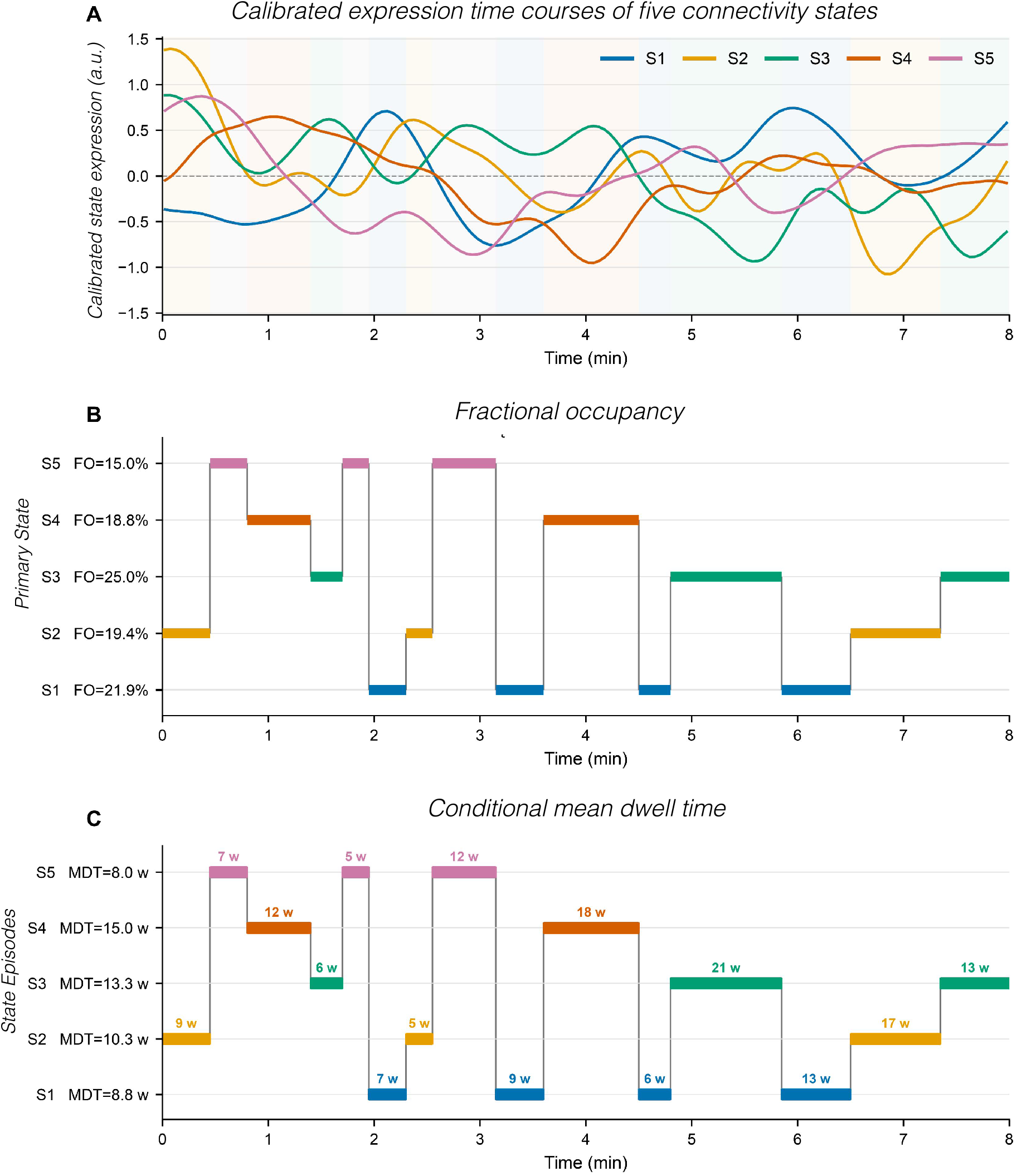
Illustration of primary-state assignment, fractional occupancy, and conditional mean dwell time at one TFM moment-frequency bin. Synthetic calibrated state-expression time courses are shown for one illustrative participant and do not represent empirical study data. (A) Continuous calibrated expression of five connectivity states. At each time point, the state with the largest absolute calibrated expression was assigned as the primary state; the light background colors indicate the resulting primary-state assignments. (B) Primary-state sequence used to calculate fractional occupancy (FO). Colored horizontal segments indicate the state assigned at each time point, and the percentage beside each state is the proportion of all time points assigned to that state. Consequently, FO sums to 100% across the five states. (C) The same primary-state sequence partitioned into contiguous episodes. Labels above the segments give episode duration in windows (w), and conditional mean dwell time (MDT) is the mean episode duration for each state. MDT is defined only for states visited by the participant. A 3-s temporal step was used to express the illustrative horizontal axis in minutes; MDT was calculated in windows.

For participant *s*, state *k*, frequency bin *f*, and rank *r*, FO was

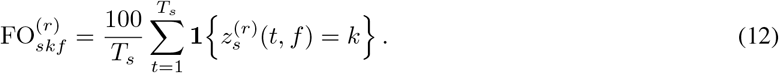

Thus, 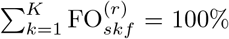 for each participant, rank, and frequency. Consecutive windows assigned to the same state were treated as episodes. If state *k* occurred in 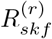 episodes with lengths 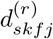, its conditional MDT was

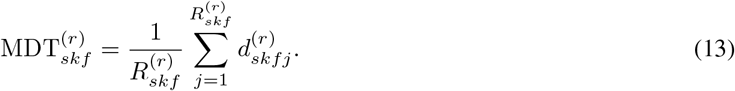

MDT was recorded in windows and was undefined when a participant never occupied the specified state at that rank and frequency. It therefore measures episode duration conditional on visitation and should not be interpreted as zero for nonvisited states.

Frequency-aggregated FO was calculated over all time–frequency observations:

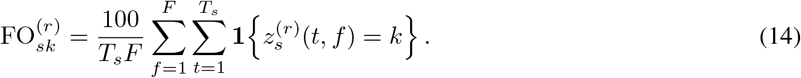

This hard ranking was used only to derive FO and MDT. The continuous calibrated amplitude of a state, 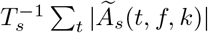, retains information from every window and is a distinct estimand.

### 2.9 Statistical Analysis of Frequency-Resolved Metrics

For each state–frequency combination, group differences were estimated with a linear model containing group and the prespecified nuisance covariates age, sex, education, and acquisition site:

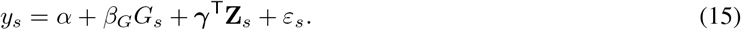

FO was analyzed on its percentage scale. Conditional MDT was log-transformed before analysis; adjusted values were exponentiated and are therefore geometric means. Figure error bars are pointwise 95% confidence intervals evaluated at the sample-average covariate values. Group inference used 50,000 Freedman–Lane permutations of the reduced-model residuals. If **ŷ**_o_ and **ê**_o_ denote fitted values and residuals from the nuisance-only model, the response in permutation *b* was

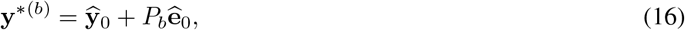

and the full model was refitted. The two-sided permutation *p* value was

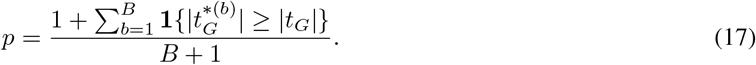

For the frequency-resolved MDT and FO panels, state–frequency comparisons with a nominal two-sided *p* < 0.05 were identified. Frequency-aggregated FO and the second-ranked sensitivity metrics were analyzed separately using two-sided Welch tests. Effect estimates for these comparisons were reported as group differences with Welch 95% confidence intervals and Hedges’ *g*.

## 3 Results

### 3.1 Derivation and Replicability of Normative Dynamic State Templates

After one-to-one matching of states across Batch A and Batch B on the basis of their edgewise Pearson correlations, all three methods recovered similar five-state structures (Figures 5 and 6). For wFNC, state-wise correlations were 0.99, 0.92, 0.98, 0.79, and 0.99 for k-means (median *r* = 0.98); 0.89, 0.93, 0.84, 0.97, and 0.90 for ICA (median *r* = 0.90); and 0.80, 0.90, 0.83, 0.94, and 0.98 for PCA (median *r* = 0.90). The states ranged from relatively segregated configurations with strong within-domain organization and opposing sensory, subcortical, and higher-cognition systems to more integrated configurations with widespread positive coupling. TFM correlations were 0.86, 0.92, 0.98, 0.89, and 0.89 for k-means (median *r* = 0.89); 0.85, 0.94, 0.88, 0.84, and 0.77 for ICA (median *r* = 0.85); and 0.68, 0.82, 0.70, 0.82, and 0.85 for PCA (median *r* = 0.82). Thus, the TFM-PCA templates were reproducible but less stable than the k-means and ICA solutions. The remaining analyses demonstrate subject-specific estimation using the ICA-derived priors.

**Figure 5:**
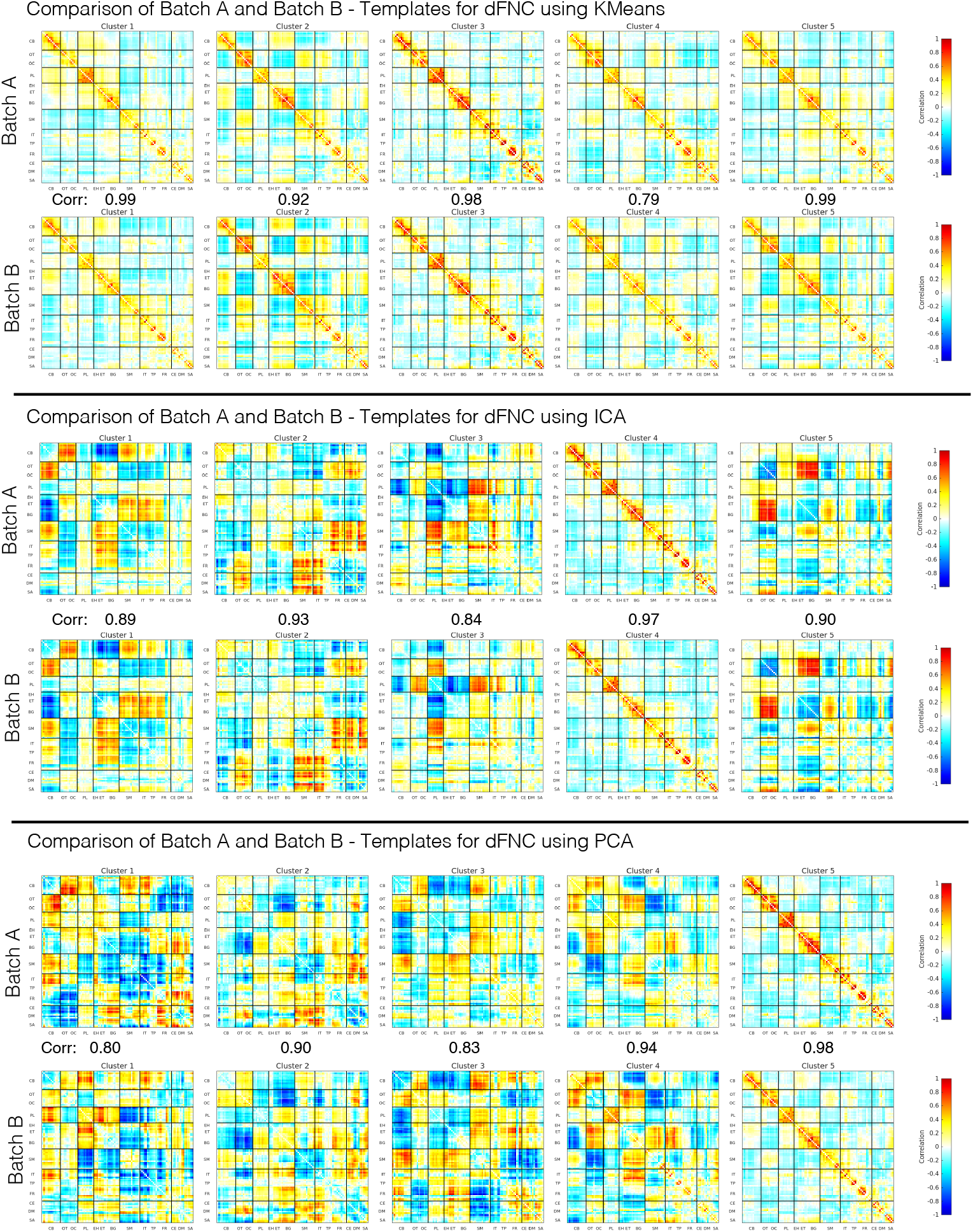
Split-sample reproducibility of the five normative windowed functional network connectivity (wFNC) state templates in the UK Biobank reference cohort. Templates were estimated independently in Batch A and Batch B using k-means, independent component analysis (ICA), and principal component analysis (PCA). Within each method, matched states are arranged in columns 1–5, with the Batch A templates shown above the corresponding Batch B templates. “Corr” denotes the Pearson correlation between each pair of matched template maps. Colors use display scale from −1 to 1, k-means panels show centroid connectivity values, whereas ICA and PCA panels show method-specific component weights rather than raw connectivity correlations. Rows and columns follow the NeuroMark 2.2 subdomain ordering in Figure 2, with black lines represent domain boundaries.

**Figure 6:**
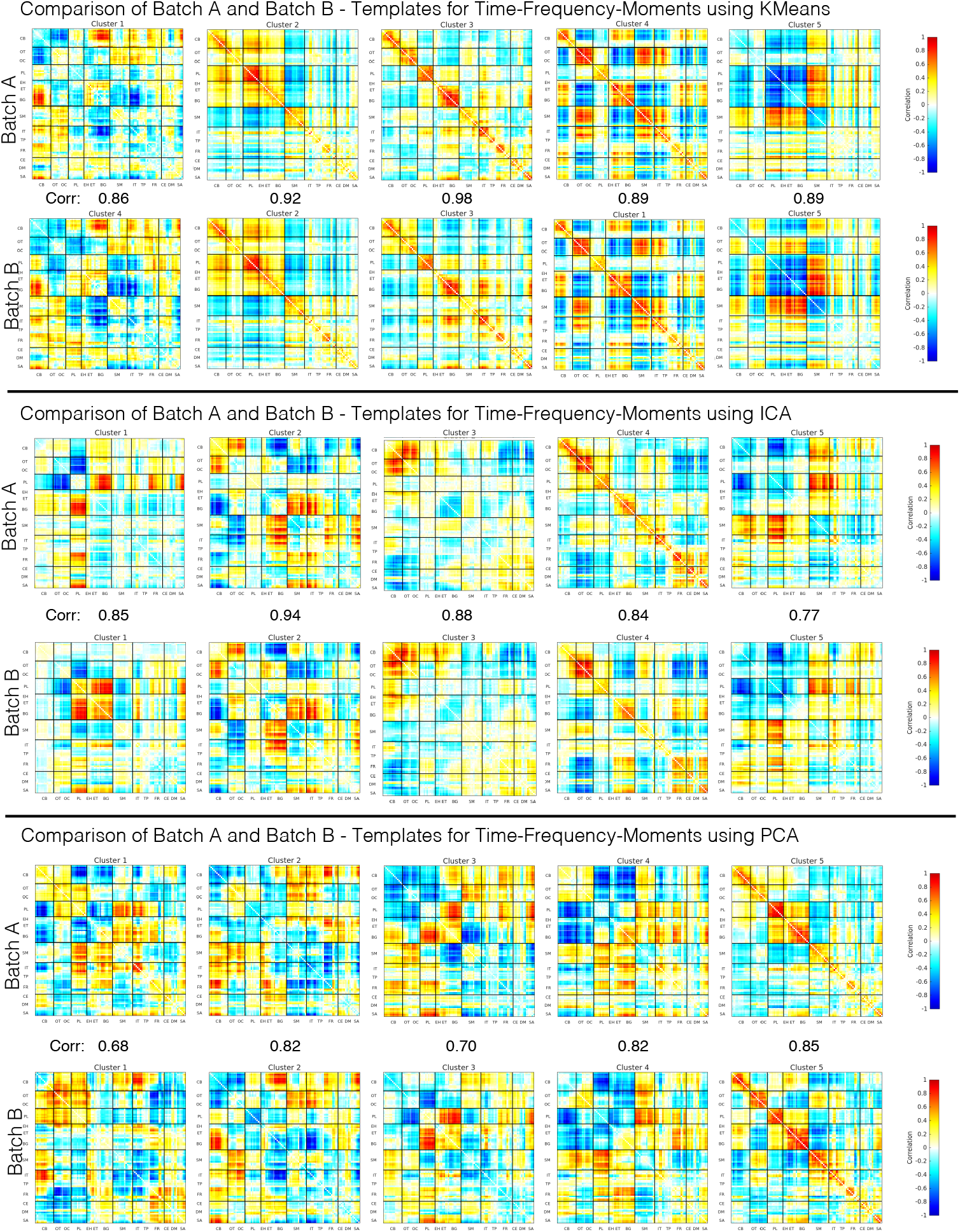
Split-sample reproducibility of the five normative time-frequency-moment (TFM) state templates in the UK Biobank reference cohort. Templates were estimated independently in Batch A and Batch B using k-means, independent component analysis (ICA), and principal component analysis (PCA). Within each method, matched states are arranged in columns 1–5, with the Batch A templates shown above the corresponding Batch B templates. “Corr” denotes the Pearson correlation between each pair of matched template maps. Colors use a common display scale from −1 to 1, k-means panels show TFM centroid values, whereas ICA and PCA panels show method-specific component weights. Rows and columns follow the NeuroMark 2.2 subdomain ordering in Figure 2, with black lines represent domain boundaries.

### 3.2 Subject-Specific State Estimation and Altered Dynamics in Schizophrenia

We next applied the UKB-derived templates to estimate subject-specific states in the independent FBIRN cohort and compared individuals with schizophrenia (SZ) and healthy controls (HC). All analyses used the five-state solution selected previously using the elbow criterion and prior research [49, 50]. The corresponding HC and SZ group-mean maps were qualitatively similar within each template-defined state, while the five states retained distinct distributed connectivity configurations (Figure 7, panels A and B). Across states, the maps differed in the sign and magnitude of within and between-domain connectivity involving visual, paralimbic, subcortical, sensorimotor, higher-cognition, and triple-network systems. These refer to the overall topology of the group-mean maps and are not tests of group differences.

**Figure 7:**
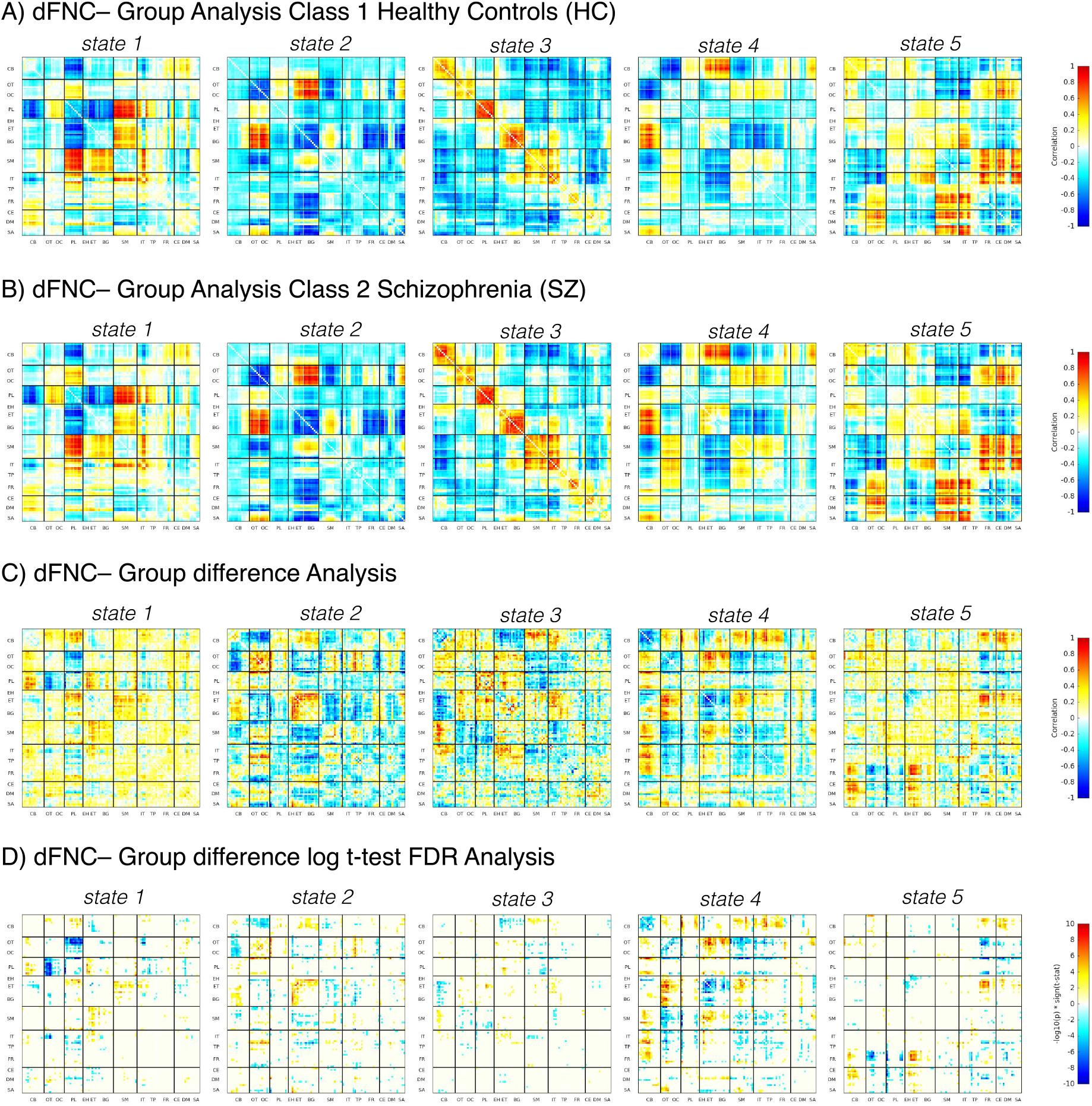
Subject-specific five-state windowed functional network connectivity patterns and case–control differences in the FBIRN cohort. (A, B) Mean state-specific connectivity matrices for healthy controls (HC) and individuals with schizophrenia (SZ), respectively. (C) Unthresholded mean group differences calculated as HC minus SZ. (D) Edges surviving false-discovery-rate correction (*q <* 0.05), encoded as sign(*t*) *×* [−log_10_(*p*)]; positive values indicate HC > SZ, negative values indicate SZ > HC, and uncolored cells denote nonsignificant effects. Matrix axes use the NeuroMark 2.2 subdomain abbreviations: CB (cerebellar); OT and OC (visual); PL (paralimbic); EH, ET, and BG (subcortical); SM (sensorimotor); IT, TP, and FR (higher cognition); and CE, DM, and SA (triple network). Full subdomain names and ICN ordering are provided in Figure 2; black lines separate subdomains.

Edgewise contrasts were state dependent rather than reflecting a uniform SZ-related shift (Figure 7, panels C and D). After FDR correction, State 4 showed spatially differences, State 3 showed comparatively sparse effects, and States 1, 2, and 5 showed more limited distributed differences. Significant edges occurred in both contrast directions and involved multiple functional domains. Because the contrasts were estimated within matched template-defined states, no post-hoc alignment of state labels was required. These findings support state-dependent, distributed dysconnectivity but do not identify a single network mechanism.

At two-sided *p <* 0.05 threshold, SZ showed a shorter conditional mean dwell time in State 2 (Figure 8, panel B). Fractional occupancy was higher in SZ for State 3 and lower for State 4 (panel C); no other state comparisons were marked at this threshold. Thus, the observed temporal differences were confined to selected primary-state summaries rather than occurring across all five states.

**Figure 8:**
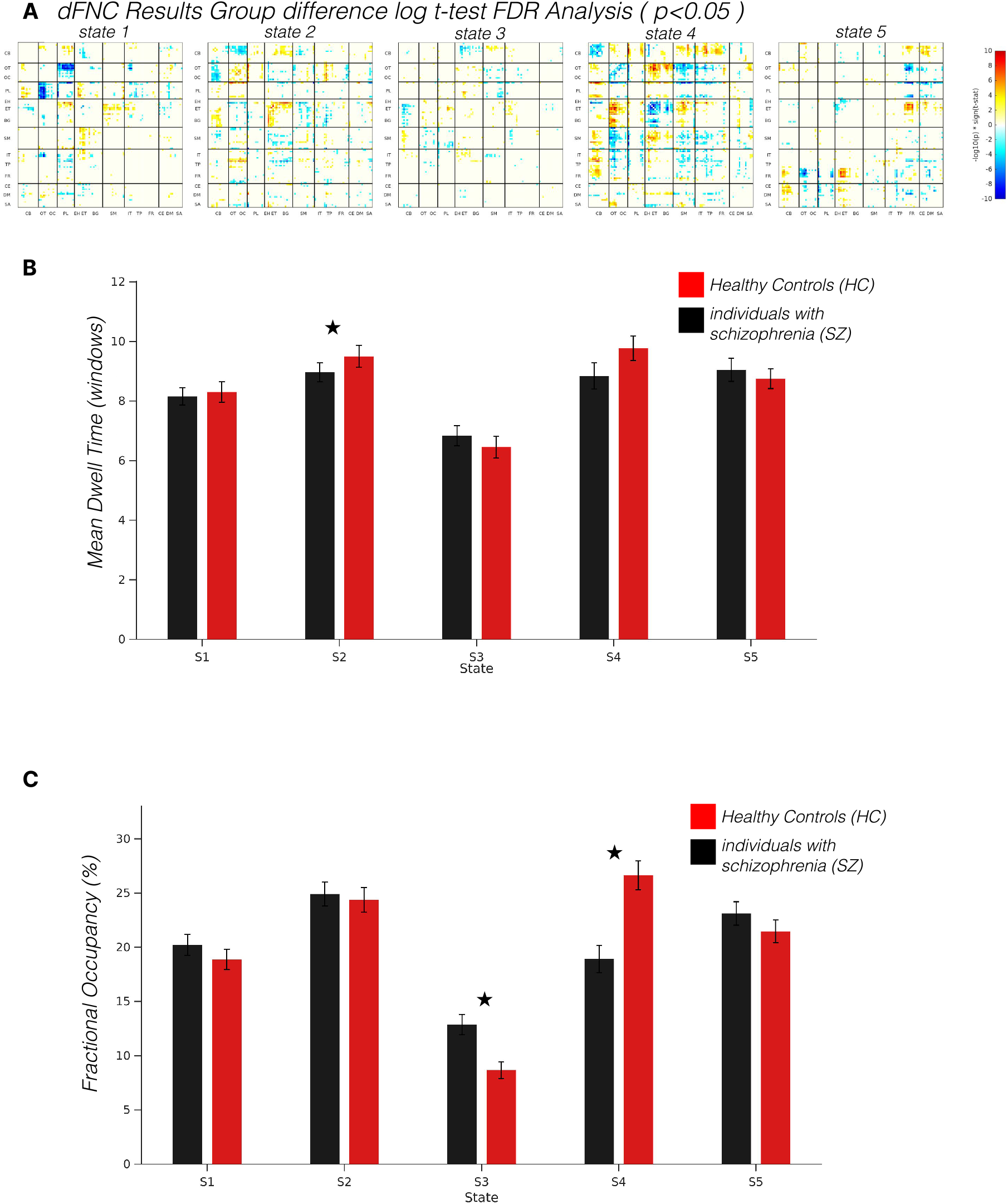
Primary-state temporal measures and thresholded edgewise differences for the wFNC analysis of the FBIRN cohort. (A) Edges surviving false-discovery-rate correction (*q <* 0.05) for the healthy-control-minus-schizophrenia contrast, encoded as sign(*t*) *×* [−log_10_(*p*)]; positive values indicate HC > SZ, negative values indicate SZ > HC, and uncolored cells denote nonsignificant effects. (B) Conditional mean dwell time for each of the five primary states. (C) Fractional occupancy for each primary state. Red bars denote HC, black bars denote SZ, and error bars show the standard error of the mean. Asterisks indicate two-sided group differences with *p <* 0.05.

### 3.3 Frequency-Aware Dynamic Dysconnectivity via Time-Frequency Moments

Incorporating TFM into NeuroMark dFNC provided an edgewise state pattern together with a frequency-resolved expression profile. The five TFM group-mean maps represented distinct configurations and were qualitatively similar between HC and SZ within corresponding states (Figure 9, panels A and B). The unthresholded contrasts are shown in panel C. After FDR correction (panel D), edgewise effects remained state dependent and involved several functional domains; effects were more numerous in States 4 and 5 and comparatively sparse in State 3. These maps do not support assigning an individual state to a single functional system.

**Figure 9:**
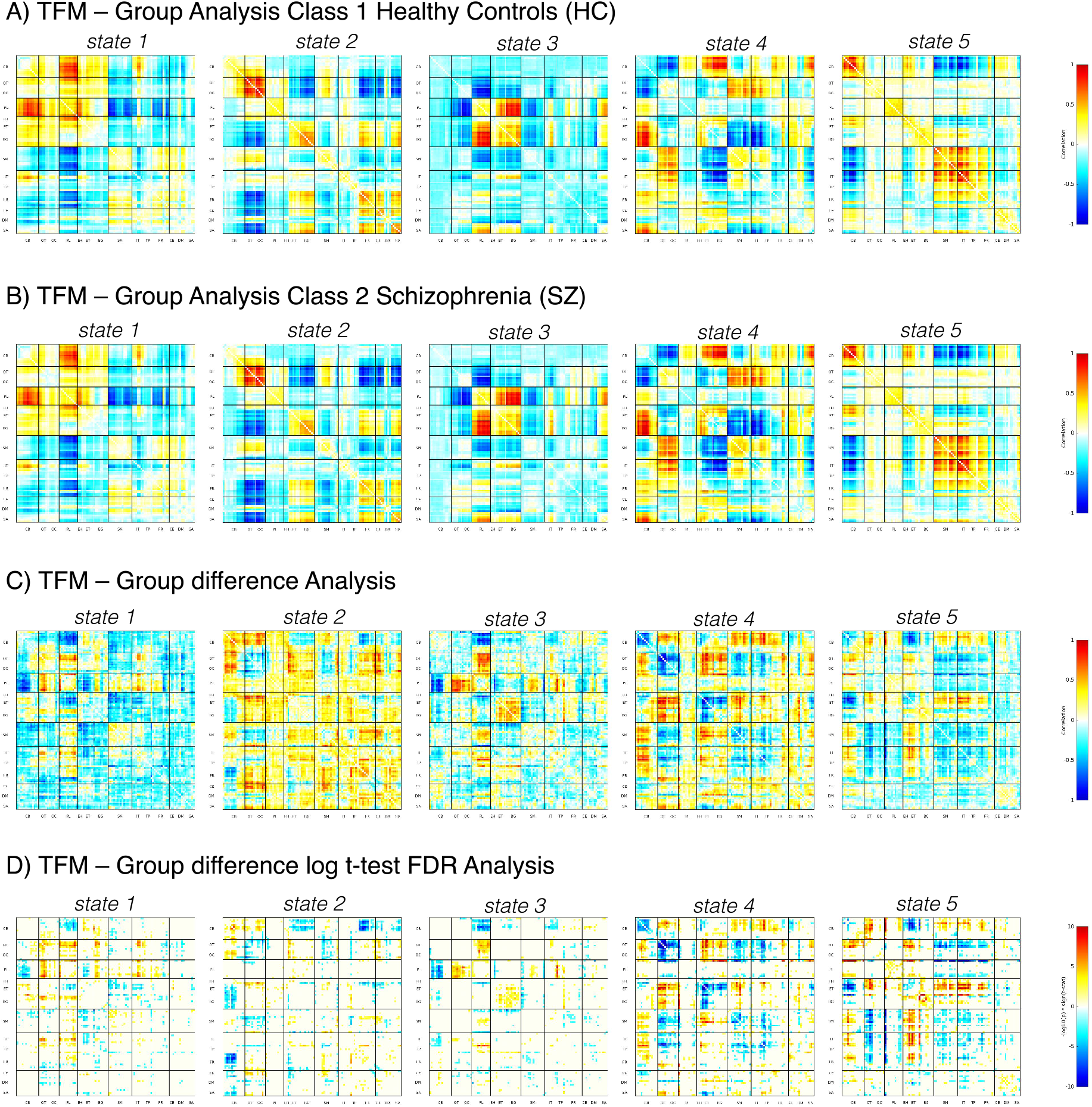
Subject-specific five-state connectivity patterns estimated with time-frequency-moment templates and case– control differences in the FBIRN cohort; each panel displays States 1–5 from left to right. (A) Mean TFM-derived state matrices for healthy controls (HC). (B) Mean TFM-derived state matrices for individuals with schizophrenia (SZ). (C) Unthresholded mean group differences calculated as HC minus SZ. (D) Edges surviving false-discovery-rate correction (*q <* 0.05), encoded as sign(*t*) *×* [−log_10_(*p*)]; positive values indicate HC > SZ, negative values indicate SZ > HC, and uncolored cells denote nonsignificant effects. Matrix rows and columns follow the NeuroMark 2.2 subdomain ordering shown in Figure 2, with black lines separating subdomains.

Frequency-resolved conditional MDT and FO are summarized in Figure 10, panels B and C, respectively. At the nominal two-sided *p <* 0.05 threshold, MDT differences were marked at one or more frequency bins in all five states. FO differences were marked for States 1, 3, and 5, whereas no bins were marked for States 2 or 4. The direction of the observed differences varied across states and frequencies. Because these state–frequency comparisons were not corrected for multiplicity, they are treated as exploratory and do not establish effects at individual frequency bins.

**Figure 10:**
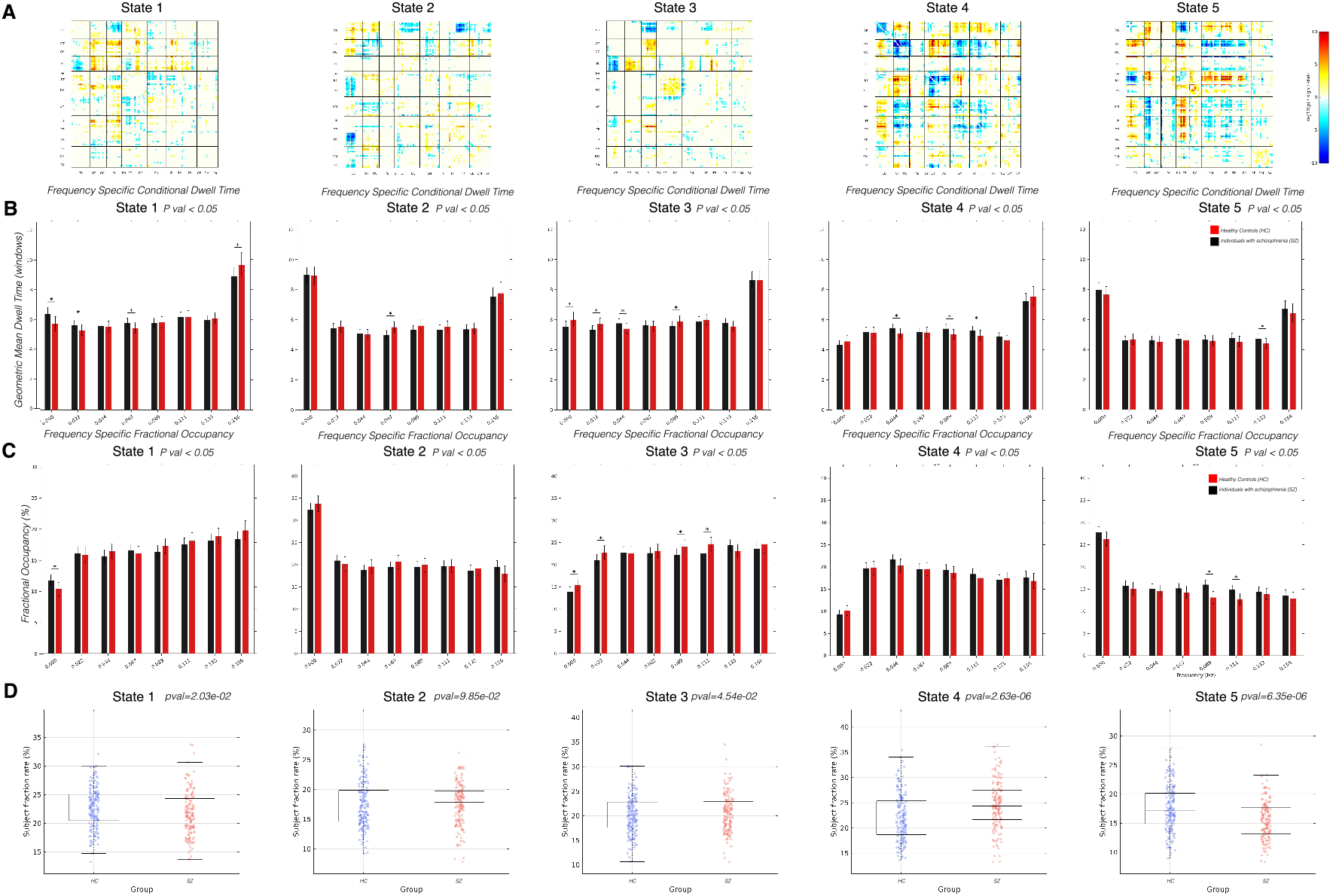
Frequency-resolved dynamics of the primary calibrated TFM state in the FBIRN cohort. At each time– frequency observation, the state with the largest absolute calibrated ICA loading was designated the primary state. (A) Edges surviving false-discovery-rate correction (*q <* 0.05) for the healthy-control-minus-schizophrenia contrast, encoded as sign(*t*) *×* [−log_10_(*p*)]; positive values indicate HC > SZ, negative values indicate SZ > HC, and uncolored cells denote nonsignificant effects. (B) Covariate-adjusted frequency-specific conditional mean dwell time, reported as geometric means in windows. (C) Covariate-adjusted frequency-specific fractional occupancy, reported as a percentage. In panels B and C, error bars are pointwise 95% confidence intervals and asterisks denote nominal two-sided *p <* 0.05. (D) Participant-level distributions of frequency-aggregated primary-state occupancy, with the nominal two-sided between-group *p* value shown for each state.

For frequency-aggregated primary-state occupancy (Figure 10, panel D), the displayed nominal tests yielded *p <* 0.05 for States 1, 3, 4, and 5, but not State 2. The smallest nominal *p* values were observed for States 4 and 5. These aggregated comparisons are reported as secondary results.

### 3.4 Cross-Disease Generalizability: Application to Alzheimer’s Disease

To assess generalizability beyond a psychiatric cohort, we applied the same UKB-derived templates to 830 cognitively normal (CN) participants and 741 participants with Alzheimer’s disease (AD) from ADNI. Subject-specific states were estimated without de novo clustering in ADNI. The corresponding CN and AD group-mean wFNC maps were qualitatively similar within each template-defined state, while distinct distributed configurations were retained across the five states (Figure 11, panels A and B).

**Figure 11:**
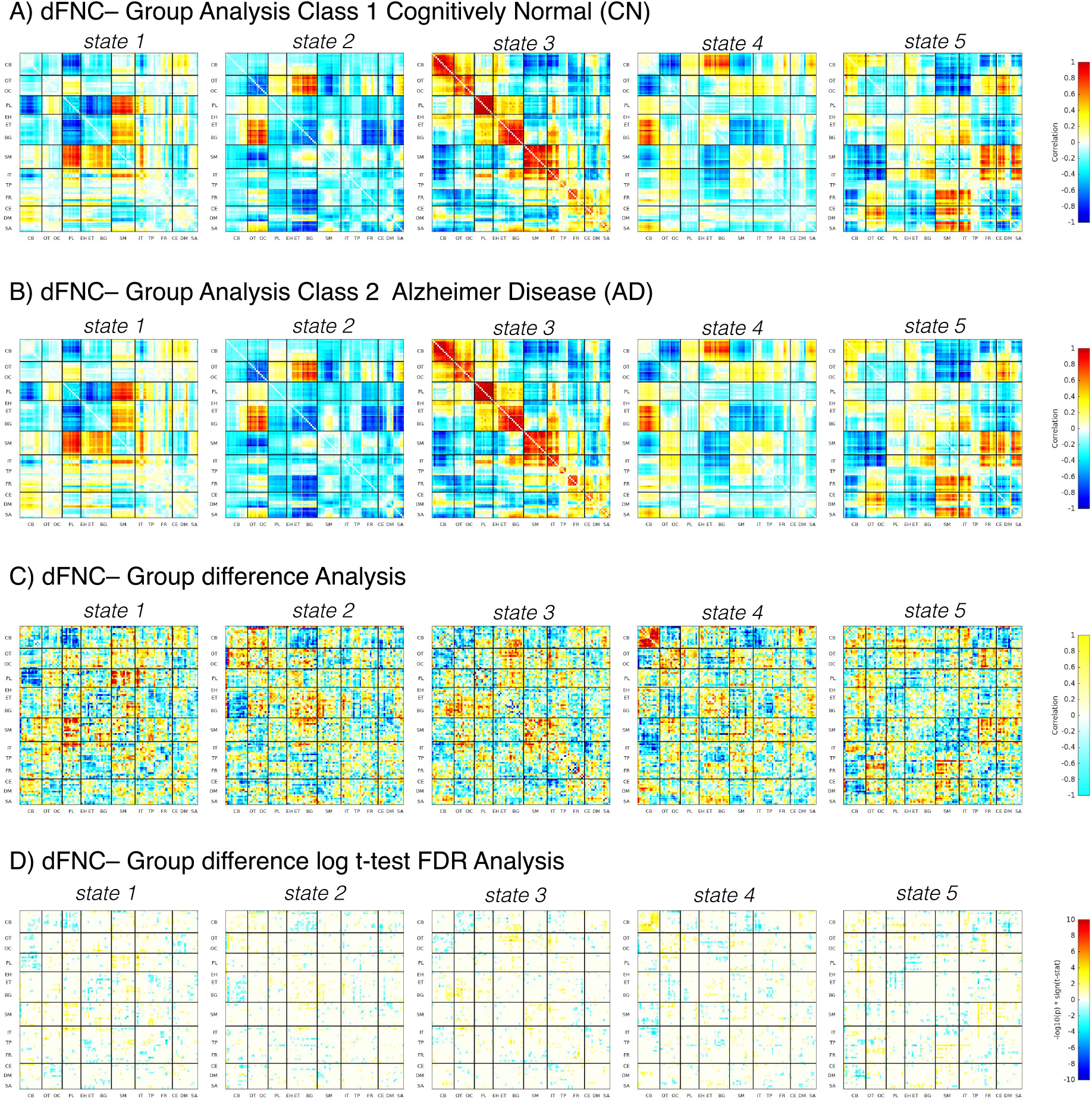
Subject-specific five-state windowed functional network connectivity patterns and diagnostic-group differences in the ADNI cohort. (A, B) Mean state-specific connectivity matrices for cognitively normal participants (CN) and participants with Alzheimer’s disease (AD), respectively. (C) Unthresholded mean group differences calculated as CN minus AD. (D) Edges surviving false-discovery-rate correction (*q <* 0.05), encoded as sign(*t*) *×* [−log_10_(*p*)]; positive values indicate CN > AD, negative values indicate AD > CN, and uncolored cells denote nonsignificant effects. Matrix rows and columns follow the NeuroMark 2.2 subdomain ordering shown in Figure 2, with black lines separating subdomains.

The unthresholded CN-minus-AD contrasts are shown in Figure 11, panel C. After FDR correction (panel D), the surviving effects were distributed across several functional domains and occurred in both contrast directions, without a uniform increase or decrease in AD. Applying the same reference templates to ADNI demonstrates transfer of the state definitions to an independent disease cohort, but it does not establish diagnostic specificity.

At the two-sided *p <* 0.05 threshold, AD showed a shorter conditional mean dwell time in State 2 (Figure 12, panel B). Fractional occupancy was lower in AD for State 2 and higher for State 4 (panel C); no other state comparisons were marked at this threshold. The observed temporal differences were therefore limited to selected primary-state summaries.

**Figure 12:**
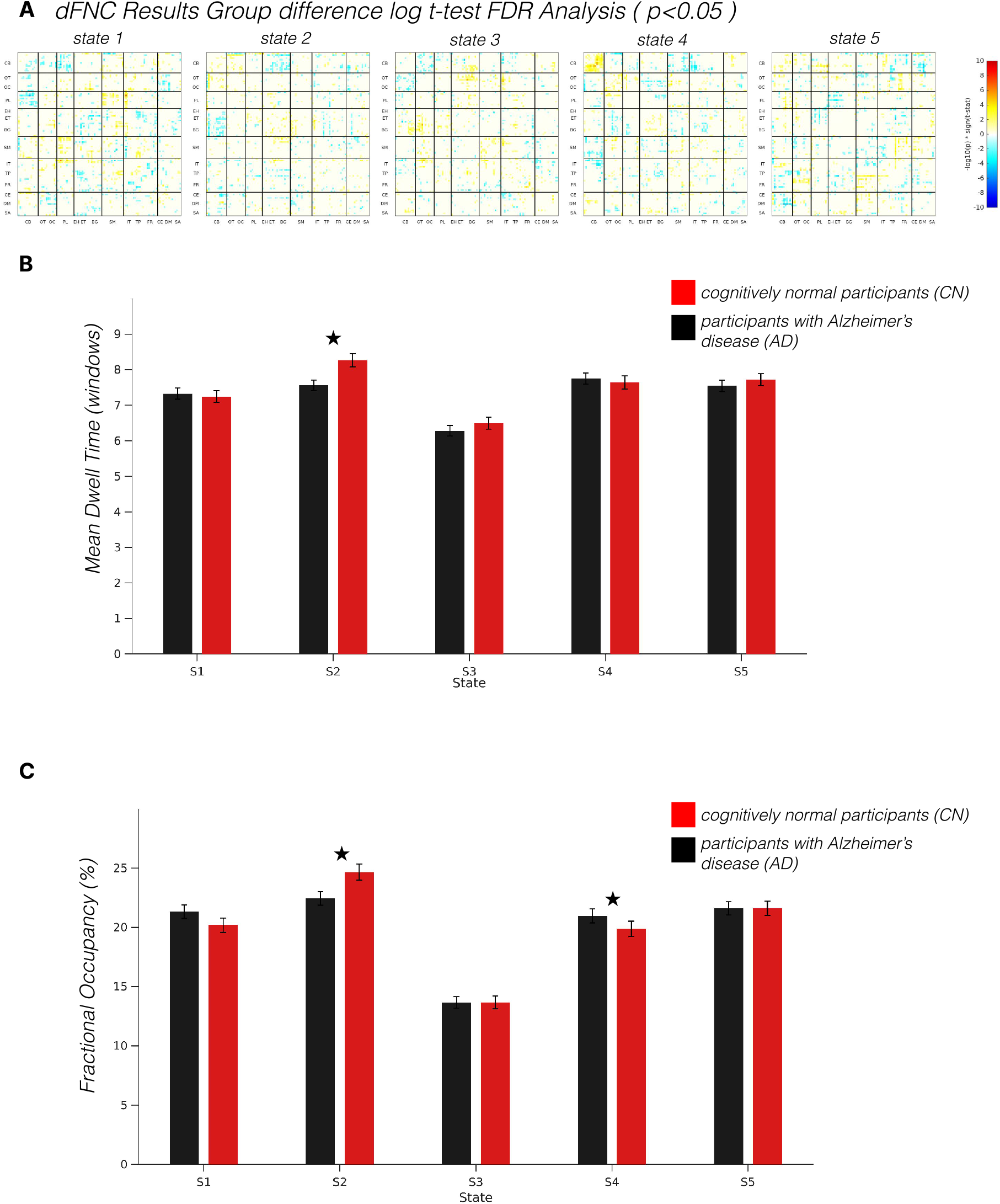
Primary-state temporal measures and thresholded edgewise differences for the wFNC analysis of the ADNI cohort. (A) Edges surviving false-discovery-rate correction (*q <* 0.05) for the cognitively-normal-minus-Alzheimer’s-disease contrast, encoded as sign(*t*) *×* [−log_10_(*p*)]; positive values indicate CN > AD, negative values indicate AD > CN, and uncolored cells denote nonsignificant effects. (B) Conditional mean dwell time for each of the five primary states. (C) Fractional occupancy for each primary state. Red bars denote CN, black bars denote AD, and error bars show the standard error of the mean. Asterisks indicate two-sided group differences with *p <* 0.05.

The TFM-derived group-mean maps also retained distinct distributed configurations and were qualitatively similar between CN and AD within corresponding states (Figure 13, panels A and B). The unthresholded contrasts are shown in panel C. After FDR correction (panel D), the surviving effects were relatively sparse, occurred in both directions, and involved edges distributed across several functional domains. Accordingly, the TFM maps were interpreted as whole-brain configurations rather than as states specific to individual networks.

**Figure 13:**
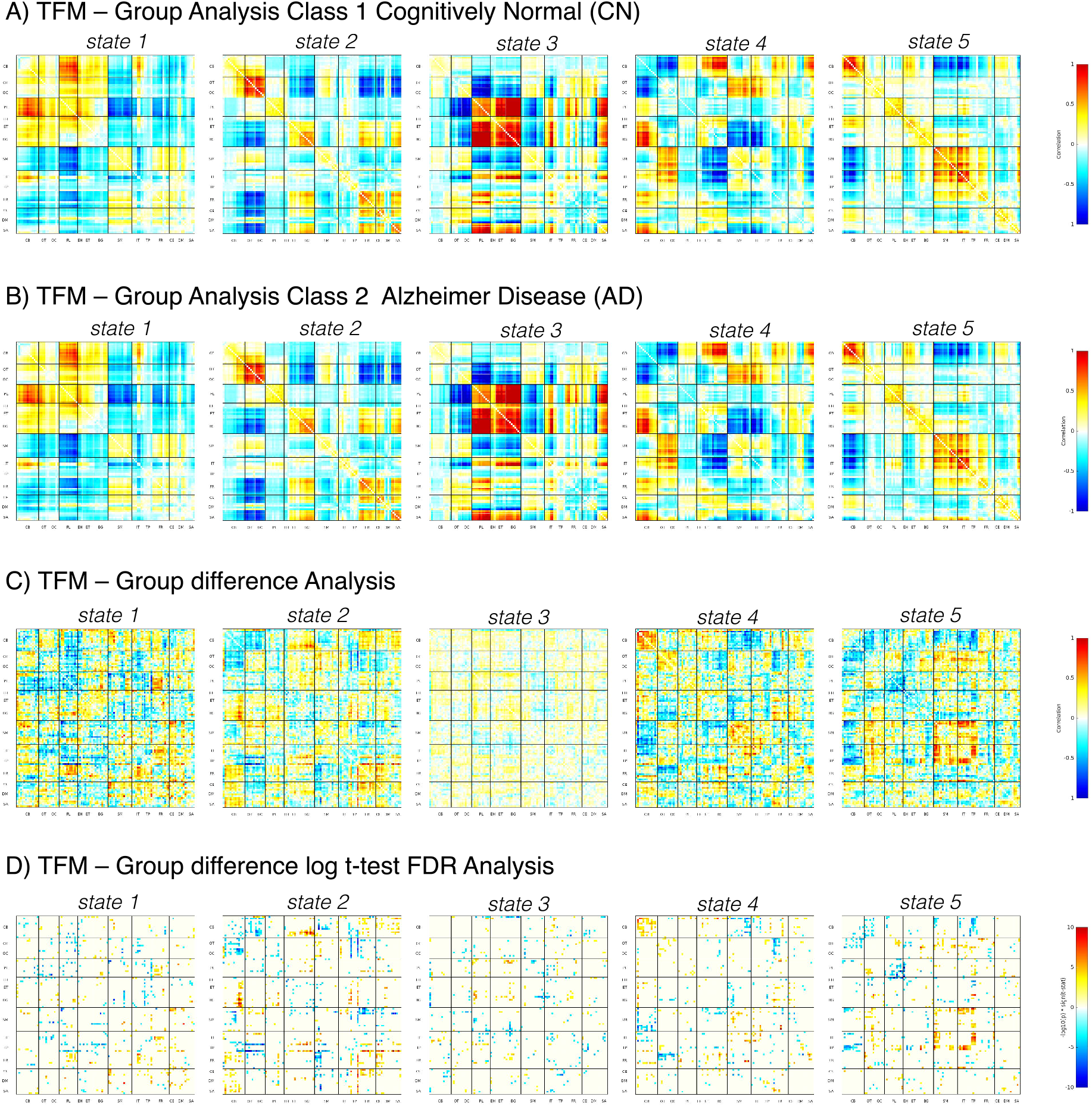
Subject-specific five-state connectivity patterns estimated with time-frequency-moment templates and diagnostic-group differences in the ADNI cohort; each panel displays States 1–5 from left to right. (A) Mean TFM-derived state matrices for cognitively normal participants (CN). (B) Mean TFM-derived state matrices for participants with Alzheimer’s disease (AD). (C) Unthresholded mean group differences calculated as CN minus AD. (D) Edges surviving false-discovery-rate correction (*q <* 0.05), encoded as sign(*t*) *×* [−log_10_(*p*)]; positive values indicate CN > AD, negative values indicate AD > CN, and uncolored cells denote nonsignificant effects. Matrix rows and columns follow the NeuroMark 2.2 subdomain ordering shown in Figure 2, with black lines separating subdomains.

Frequency conditional MDT and FO are in Figure 14, panels B and C, respectively. At two-sided *p <* 0.05 threshold, MDT differences were marked at one or more bins for States 2-5, with no marked bin for State 1. FO differences were marked for all five states; State 5 contained the largest number of marked bins. Directions varied across states and frequencies. Because these state-frequency comparisons were not corrected for multiplicity, they are treated as exploratory and do not establish effects at individual frequency bins.

**Figure 14:**
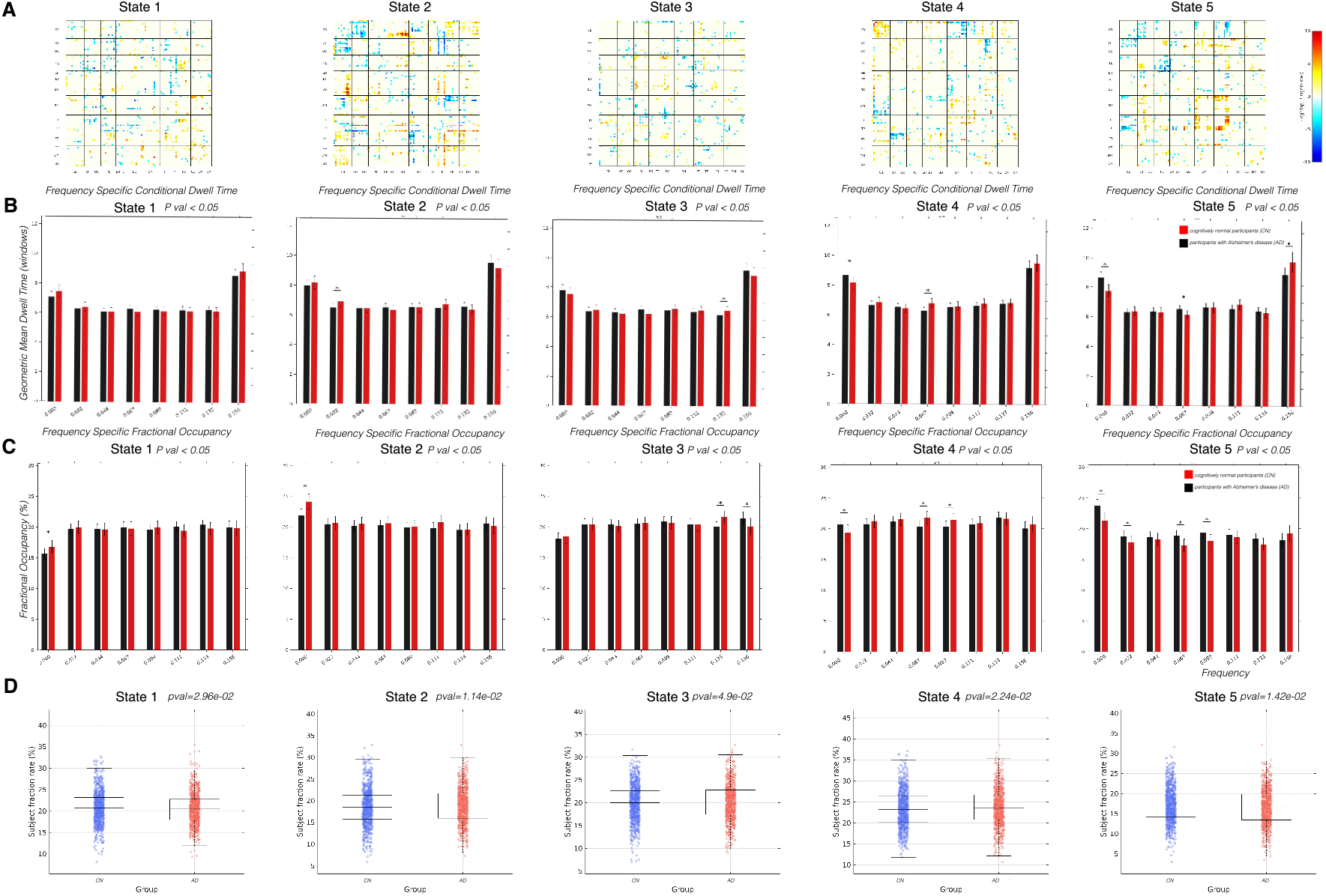
Frequency dynamics of the calibrated TFM state in the ADNI cohort. At each time-frequency observation, the state with the largest absolute calibrated ICA loading was designated the primary state. (A) Edges surviving false-discovery-rate correction (*q <* 0.05) for the cognitively-normal-minus-Alzheimer’s-disease contrast, encoded as sign(*t*) *×* [−log_10_(*p*)]; positive values indicate CN > AD, negative values indicate AD > CN, and uncolored cells denote nonsignificant effects. (B) Covariate-adjusted frequency-specific conditional mean dwell time, reported as geometric means in windows. (C) Covariate-adjusted frequency-specific fractional occupancy, reported as a percentage. In panels B and C, error bars are pointwise 95% confidence intervals and asterisks denote nominal two-sided *p <* 0.05. (D) Participant-level distributions of frequency-aggregated primary-state occupancy, with the nominal two-sided between-group *p* value shown for each state.

Panel D of Figure 14 shows the participant-level distributions of frequency-aggregated primary-state occupancy. The nominal tests yielded *p <* 0.05 for all five states, but the group distributions overlapped substantially. These results therefore indicate small group shifts in aggregated occupancy and are treated as secondary rather than evidence of a broad redistribution of state expression.

## 4 Discussion

dFNC has provided important evidence that large-scale brain networks fluctuate among recurring connectivity configurations rather than remaining static over the course of resting-state fMRI acquisition [10, 11, 13]. However, a major barrier to the clinical translation of dFNC has been the limited reproducibility of data-driven state definitions. In conventional pipelines, dynamic states are typically derived separately for each dataset using clustering or decomposition methods, making the resulting states dependent on sample composition, acquisition characteristics, preprocessing choices, and model parameters such as the number of clusters or window length [23, 15, 39, 51]. This creates a fundamental difficulty for validation studies: a state identified in one cohort may not correspond directly to a state identified in another cohort, and newly acquired subjects cannot be easily mapped onto an established state space. The present work addresses this limitation by introducing NeuroMark dFNC, a template-guided and fully automated framework for estimating subject-specific dynamic connectivity states using reproducible normative priors.

The central contribution of NeuroMark dFNC is the extension of prior-guided modeling from spatial network estimation to dynamic state estimation. Prior NeuroMark work has demonstrated that spatially constrained ICA can provide reproducible intrinsic connectivity network estimates across subjects, datasets, and disorders by using reliable spatial network templates as priors [5, 6, 36]. Here, we apply the same general principle to the dynamic connectivity domain: instead of allowing each dataset to define its own state space, we construct dFNC state templates from a large reference cohort and use those templates to guide subject-level estimation in independent datasets. This approach creates a common coordinate system for dynamic brain states, enabling direct comparison across individuals and cohorts without post-hoc state matching. Importantly, the split-half analyses in the UK Biobank dataset demonstrated that the derived templates were highly stable across independent batches, supporting their use as reproducible priors rather than dataset-specific artifacts.

An important feature of the ICA formulation is that state expression remains continuous rather than mutually exclusive. The discrete labels used for FO and MDT are therefore summaries of an underlying soft state representation, allowing overlapping state expression while retaining familiar occupancy and dwell-time measures.

The schizophrenia findings illustrate the clinical value of this approach. Using UK Biobank-derived templates in the independent FBIRN cohort, NeuroMark dFNC identified state-specific dysconnectivity involving default mode, sensorimotor, visual, higher cognitive, and subcortical systems. These findings are consistent with the long-standing dysconnectivity hypothesis of schizophrenia, which proposes that psychotic symptoms and cognitive impairment arise from abnormal integration and segregation among distributed brain networks [52, 53, 54]. In particular, the attenuation of default mode–sensorimotor and default mode–visual antagonism suggests reduced functional segregation between internally oriented and externally oriented systems. Such impaired segregation may contribute to clinical features of schizophrenia including altered self-referential processing, impaired sensory filtering, and difficulty distinguishing internally generated from externally driven information.

The observed involvement of subcortical and thalamocortical connections is especially clinically meaningful. The thalamus plays a key role in coordinating cortical communication, sensory gating, attention, and cognitive control. Abnormal thalamocortical connectivity has been repeatedly reported in schizophrenia [55, 56] and is thought to contribute to disrupted information flow between sensory, association, and executive systems [57, 58]. In the present study, schizophrenia was associated with abnormal expression of a globally hyperconnected state and altered subcortical-cortical connectivity. This pattern may reflect a failure of dynamic gating mechanisms, in which the brain becomes less able to flexibly regulate communication between subcortical hubs [59] and distributed cortical networks. Clinically, such disruptions and connectivity patterns may be relevant to hallucinations, disorganized thought, attentional instability, and cognitive dysfunction [60, 61, 62, 63].

Beyond edgewise differences in connectivity, the frequency-specific temporal analysis identified HC–SZ differences in conditional mean dwell time and fractional occupancy at selected state–frequency combinations. The directions of these effects were heterogeneous across neighboring frequencies, so the findings do not support a single global increase or decrease in state persistence. Instead, they indicate that schizophrenia-related temporal alterations depend on both state identity and moment frequency. These metrics may provide useful intermediate phenotypes, although future studies must establish their relationships with symptoms, cognition, medication exposure, and longitudinal outcomes.

A second important contribution of this work is the incorporation of time-frequency moment templates into dFNC estimation. Standard wFNC captures temporal variation in connectivity but does not explicitly resolve the moment-frequency structure of pairwise interactions. The TFM analysis paired edgewise state patterns with frequency-specific temporal metrics, showing that diagnostic-group effects varied across states and moment-frequency bins. This frequency-resolved dimension may provide complementary information by identifying both the network topology of an altered state and the frequencies at which its expression differs between groups. Importantly, TFM frequencies describe variation in statistical-moment space and should not be interpreted as electrophysiological frequency bands.

Application to the ADNI dataset further demonstrated the cross-disease generalizability of NeuroMark dFNC. Alzheimer’s disease is known to affect large-scale functional networks, especially the default mode network, posterior cingulate/precuneus, medial temporal, and higher-order associative systems [64, 65, 66]. These regions overlap strongly with the topography of amyloid and tau vulnerability and are central to episodic memory, self-referential processing, and cognitive integration. In the present study, the same UK Biobank-derived templates that captured schizophrenia-related dysconnectivity also detected Alzheimer’s-related alterations, including state-specific edgewise differences and modest frequency-dependent differences in conditional dwell time and fractional occupancy. Because the temporal effects varied across states and frequencies, they should not be interpreted as evidence for uniformly preserved or impaired switching, or as proof of disorder-specific mechanisms.

The Alzheimer’s disease findings have important clinical implications. Functional connectivity changes may emerge before severe structural atrophy and may provide information about network-level dysfunction that is not captured by anatomy alone [64, 67]. A template-guided dFNC framework could therefore support more standardized evaluation of dynamic connectivity alterations along the Alzheimer’s disease continuum, including cognitively normal individuals at elevated risk, mild cognitive impairment, and clinically diagnosed dementia. Frequency-specific shifts in fractional occupancy or conditional dwell time may reflect altered allocation of dynamic state expression, but their biological and clinical meaning cannot be established from the present group comparison alone. Future work linking NeuroMark dFNC measures to amyloid, tau, neurodegeneration markers, cognitive decline, and longitudinal conversion will be essential for determining whether these dynamic features provide clinically actionable biomarkers.

Taken together, the schizophrenia and Alzheimer’s disease applications highlight the broader value of NeuroMark dFNC as a transdiagnostic framework. Rather than defining a new set of states for each disorder, the same normative state templates were applied across independent cohorts and disease domains. This design makes it possible to ask whether different disorders alter the same dynamic states in different ways, or whether they preferentially affect distinct aspects of the dynamic repertoire. Such a framework is especially important for clinical translation because a biomarker must be stable enough to generalize across studies while remaining sensitive enough to capture disease-specific variation. NeuroMark dFNC directly addresses this balance by combining normative priors with subject-specific estimation.

This work also has implications for future individualized neuroimaging. Because the proposed framework maps each subject into a common dynamic state space, it can support subject-level metrics such as state expression, fractional occupancy, dwell time, and state-specific connectivity deviations. These features may be useful for predictive modeling, patient stratification, and longitudinal monitoring. For example, the state- and frequency-specific measures identified here could be tested for associations with symptom severity or cognitive impairment in schizophrenia and with disease stage or progression in Alzheimer’s disease. Importantly, the availability of shared templates would allow different laboratories to estimate comparable dynamic features, facilitating replication and meta-analysis across cohorts.

Several limitations should be considered. First, the wFNC representation remains sensitive to window length, sampling rate, preprocessing, and noise characteristics. The same 40-TR window also corresponds to different durations across datasets. Robustness should be evaluated across window parameters and against non-windowed approaches such as hidden Markov models, co-activation patterns, and point-process methods [29, 32]. Second, FO and conditional MDT discard part of the continuous state-expression information. MDT is also undefined for participants who do not visit a given state, so an apparent group difference may reflect episode duration, differential visitation, or both. Reporting visitation probability and continuous calibrated amplitude would help separate these mechanisms. Third, the present analyses are group comparisons; links with symptoms, cognition, medication, disease duration, biomarkers, and longitudinal outcomes remain to be tested.

Another limitation is that the normative templates were derived from UK Biobank, a large and high-quality cohort, but one that has its own demographic and acquisition characteristics. Although the successful application to FBIRN and ADNI supports generalizability, future studies should evaluate template performance across broader age ranges, scanner protocols, ethnic backgrounds, and clinical populations. It will also be important to examine whether disease-specific or age-specific template sets provide additional sensitivity beyond a single normative template set. Finally, while the current study demonstrates promising group-level differences, clinical translation will require rigorous test-retest reliability, site harmonization, external validation, and prediction of individual outcomes.

In summary, NeuroMark dFNC provides a reproducible and scalable framework for estimating subject-specific dynamic connectivity states from resting-state fMRI. By treating dynamic states as transferable templates rather than dataset-specific clustering outputs, the proposed pipeline improves cross-cohort comparability while preserving sensitivity to individual and clinical variation. The results demonstrate that stable normative dFNC templates can be applied to independent schizophrenia and Alzheimer’s disease cohorts to reveal biologically plausible, disease-relevant alterations in both connectivity patterns and temporal state dynamics. By sharing the derived templates with the community, this work provides a step toward standardized dynamic connectomics and supports future efforts to develop reliable, clinically meaningful dFNC biomarkers. The group-level template files are available through Supplemental Data to support independent evaluation and reuse.

## 5 Conclusion

This study introduced NeuroMark dFNC, a fully automated, template-guided framework for estimating subject-specific dynamic functional connectivity states from resting-state fMRI. By using reproducible state templates derived from a large reference cohort, the proposed approach addresses a key limitation of conventional data-driven dFNC pipelines: the difficulty of replicating brain states across datasets and applying them consistently to new subjects. Across independent schizophrenia and Alzheimer’s disease cohorts, NeuroMark dFNC captured group differences in state-specific connectivity patterns and frequency-resolved temporal measures. The schizophrenia results highlighted disrupted network segregation, altered subcortical–cortical connectivity, and state- and frequency-dependent temporal differences, while the Alzheimer’s disease findings showed distinct alterations involving higher-cognitive and default-mode-related systems together with modest frequency-dependent temporal effects. The TFM-derived templates and their frequency-resolved expression profiles provided information complementary to the edgewise state maps. Overall, NeuroMark dFNC provides a reproducible and scalable framework for standardized dynamic connectomics. By treating dynamic states as transferable templates rather than dataset-specific clustering outputs, this approach supports cross-cohort comparison, individualized brain-state estimation, and future development of clinically relevant dFNC biomarkers.

## Supplemental Data

Group-level wFNC and TFM dynamic-state templates are available through Supplemental Data. The shared files contain aggregate template patterns; individual participant data are not distributed because of cohort data-sharing restrictions.

## Acknowledgments

This work was supported by the National Institutes of Health (grant numbers R01EB006841 and R01MH123610) and the National Science Foundation (grant numbers 2316421 and 2112455).

Data used in this study were obtained in part from the Function Biomedical Informatics Research Network (FBIRN with papers [40, 44]), the UK Biobank, and the Alzheimer’s Disease Neuroimaging Initiative (ADNI with papers [45]). This research was conducted using the UK Biobank Resource under Application Number 34175. Data collection and sharing for the Alzheimer’s Disease Neuroimaging Initiative (ADNI) is funded by the National Institute on Aging (National Institutes of Health Grant U19AG024904). The grantee organization is the Northern California Institute for Research and Education. In the past, ADNI has also received funding from the National Institute of Biomedical Imaging and Bioengineering, the Canadian Institutes of Health Research, and private-sector contributions through the Foundation for the National Institutes of Health (FNIH).

## Notes

https://trendscenter.org/data/

### Competing Interest Statement

The authors have declared no competing interest.

## References

[1] B. Biswal, F. Z. Yetkin, V. M. Haughton, and J. S. Hyde. Functional connectivity in the motor cortex of resting human brain using echo-planar MRI. Magnetic Resonance in Medicine, 34(4):537–541, 1995.

[2] C. F. Beckmann, M. DeLuca, J. T. Devlin, and S. M. Smith. Investigations into resting-state connectivity using independent component analysis. Philosophical Transactions of the Royal Society B: Biological Sciences, 360(1457):1001–1013, 2005.

[3] V. D. Calhoun, T. Adali, G. D. Pearlson, and J. J. Pekar. A method for making group inferences from functional MRI data using independent component analysis. Human Brain Mapping, 14(3):140–151, 2001.

[4] Masoud Seraji, Sarah Shultz, Qiang Li, Zening Fu, Vince D Calhoun, and Armin Iraji. Spatial development of brain networks during the first six postnatal months. Communications biology, 8(1):1514, 2025.

[5] Y. Du, Z. Fu, J. Sui, S. Gao, Y. Xing, D. Lin, M. Salman, A. Abrol, M. A. Rahaman, J. Chen, J. A. Turner, G. Pearlson, A. Iraji, T. Eichele, Q. Yu, J. Liu, and V. D. Calhoun. NeuroMark: An automated and adaptive ICA pipeline for measuring brain function from fMRI. NeuroImage, 222:117267, 2020.

[6] Z. Fu, Y. Tu, C. G. Abdallah, J. H. Krystal, V. D. Calhoun, and Y. Du. Test-retest reliability of spatial ICA and NeuroMark templates for fMRI. NeuroImage, 243:118548, 2021.

[7] Zening Fu, Ishaan Batta, Lei Wu, Anees Abrol, Oktay Agcaoglu, Mustafa S Salman, Yuhui Du, Armin Iraji, Sarah Shultz, Jing Sui, et al. Searching reproducible brain features using NeuroMark: templates for different age populations and imaging modalities. Neuroimage, 292:120617, 2024.

[8] Armin Iraji, Zening Fu, Ashkan Faghiri, Marlena Duda, Jaiyu Chen, Srinivas Rachakonda, Thomas DeRamus, Peter Kochunov, Bhim M Adhikari, Aysenil Belger, et al. Identifying canonical and replicable multi-scale intrinsic connectivity networks in 100k+ resting-state fmri datasets. Human Brain Mapping, 44(17):5729–5748, 2023.

[9] Masoud Seraji, Sarah Shultz, Qiang Li, Liang Ma, Zening Fu, and Vince D Calhoun. Decoding brain dynamics via cross-network non-linear trajectories. In ICASSP 2026-2026 IEEE International Conference on Acoustics, Speech and Signal Processing (ICASSP), pages 7962–7966. IEEE, 2026.

[10] C. Chang and G. H. Glover. Time–frequency dynamics of resting-state brain connectivity measured with fMRI. NeuroImage, 50(1):81–98, 2010.

[11] R. M. Hutchison, T. Womelsdorf, E. A. Allen, P. A. Bandettini, V. D. Calhoun, M. Corbetta, S. Della Penna, J. H. Duyn, G. H. Glover, J. Gonzalez-Castillo, D. A. Handwerker, S. Keilholz, V. Kiviniemi, D. A. Leopold, F. de Pasquale, O. Sporns, M. Walter, and C. Chang. Dynamic functional connectivity: promise, issues, and interpretations. NeuroImage, 80:360–378, 2013.

[12] Ünal Sakoğlu, Godfrey D Pearlson, Kent A Kiehl, Y Michelle Wang, Andrew M Michael, and Vince D Calhoun. A method for evaluating dynamic functional network connectivity and task-modulation: application to schizophrenia. Magnetic Resonance Materials in Physics, Biology and Medicine, 23(5–6):351–366, 2010.

[13] E. A. Allen, E. Damaraju, S. M. Plis, E. B. Erhardt, T. Eichele, and V. D. Calhoun. Tracking whole-brain connectivity dynamics in the resting state. Cerebral Cortex, 24(3):663–676, 2014.

[14] Ashkan Faghiri, Armin Iraji, Noah Lewis, Kun Yang, Koko Ishizuka, Akira Sawa, Tülay Adali, and Vince Calhoun. Going from lines to triangles: A formulation for time-frequency moments of time-series with application to study fmri. In 2022 IEEE 14th Image, Video, and Multidimensional Signal Processing Workshop (IVMSP), pages 1–5, 2022.

[15] S. Shakil, C. H. Lee, and S. D. Keilholz. Evaluation of sliding window correlation performance for characterizing dynamic functional connectivity and brain states. NeuroImage, 133:111–128, 2016.

[16] E. Damaraju, E. A. Allen, A. Belger, J. M. Ford, S. McEwen, D. H. Mathalon, B. A. Mueller, G. D. Pearlson, S. G. Potkin, A. Preda, J. A. Turner, J. G. Vaidya, T. G. M. van Erp, and V. D. Calhoun. Dynamic functional connectivity analysis reveals transient states of dysconnectivity in schizophrenia. NeuroImage: Clinical, 5:298–308, 2014.

[17] B. Rashid, E. Damaraju, G. D. Pearlson, and V. D. Calhoun. Dynamic functional network connectivity in the resting brain. NeuroImage, 140:8–23, 2016.

[18] Yuhui Du, Susanna L Fryer, Zening Fu, Dongdong Lin, Jing Sui, Jiayu Chen, Eswar Damaraju, Eva Mennigen, Barbara Stuart, Rachel L Loewy, Daniel H Mathalon, and Vince D Calhoun. Dynamic functional connectivity impairments in early schizophrenia and clinical high-risk for psychosis. NeuroImage, 180(Pt B):632–645, 2017.

[19] Yuhui Du, Zening Fu, Jing Sui, Shuang Gao, Ying Xing, Dongdong Lin, Mustafa Salman, Anees Abrol, Md Abdur Rahaman, Jiayu Chen, et al. Neuromark: an automated and adaptive ica based pipeline to identify reproducible fmri markers of brain disorders. NeuroImage: Clinical, 28:102375, 2020.

[20] Kaustubh Supekar, Vinod Menon, Daniel Rubin, Mark Musen, and Michael D Greicius. Network analysis of intrinsic functional brain connectivity in alzheimer’s disease. PLoS Computational Biology, 4(6):e1000100, 2008.

[21] David T Jones, Prashanthi Vemuri, Matthew C Murphy, Jeffrey L Gunter, Matthew L Senjem, Mary M Machulda, Scott A Przybelski, Brian E Gregg, Kejal Kantarci, David S Knopman, Bradley F Boeve, Ronald C Petersen, and Clifford R Jack. Non-stationarity in the “resting brain’s” modular architecture. PLoS ONE, 7(6):e39731, 2012.

[22] Sophie Dautricourt, Julie Gonneaud, Brigitte Landeau, Vince D Calhoun, Robin de Flores, Géraldine Poisnel, Salma Bougacha, Valentin Ourry, Edelweiss Touron, Elizabeth Kuhn, et al. Dynamic functional connectivity patterns associated with dementia risk. Alzheimer’s Research & Therapy, 14(1):72, 2022.

[23] Rikkert Hindriks, Mohit H Adhikari, Yuko Murayama, Marco Ganzetti, Dante Mantini, Nikos K Logothetis, and Gustavo Deco. Can sliding-window correlations reveal dynamic functional connectivity in resting-state fmri? NeuroImage, 127:242–256, 2016.

[24] Sadia Shakil, Chin-Hui Lee, and Shella D Keilholz. Parametric dependencies of sliding window correlation. IEEE Transactions on Biomedical Engineering, 65(2):254–263, 2018.

[25] Maziar Yaesoubi, Tü lay Adali, and Vince D Calhoun. A window-less approach for capturing time-varying connectivity in fmri data reveals the presence of states with variable rates of change. Human Brain Mapping, 39(4):1626–1636, 2018.

[26] Enrico Glerean, Juha Salmi, Juha M Lahnakoski, Iiro P Jääskeläinen, and Mikko Sams. Functional magnetic resonance imaging phase synchronization as a measure of dynamic functional connectivity. Brain Connectivity, 2(2):91–101, 2012.

[27] Maziar Yaesoubi, Tülay Adali, and Vince D. Calhoun. A window-less approach for capturing time-varying connectivity in fmri data reveals the presence of states with variable rates of change. Human Brain Mapping, 39(4):1626–1636, 2018.

[28] Sir-Lord Wiafe, Najme Soleimani, and Vince D. Calhoun. Dynamic co-modulation (DyCoM): A unified operator framework for dynamic connectivity in neuroimaging. bioRxiv, 2026.

[29] X. Liu and J. H. Duyn. Time-varying functional network information extracted from brief instances of spontaneous brain activity. Proceedings of the National Academy of Sciences, 110(11):4392–4397, 2013.

[30] E. Tagliazucchi, F. Von Wegner, A. Morzelewski, V. Brodbeck, and H. Laufs. Dynamic BOLD functional connectivity in humans and its electrophysiological correlates. Frontiers in Human Neuroscience, 6:339, 2012.

[31] Diego Vidaurre, Stephen M Smith, and Mark W Woolrich. Brain network dynamics are hierarchically organized in time. Proceedings of the National Academy of Sciences, 114(48):12827–12832, 2017.

[32] Diego Vidaurre, Laurence T Hunt, Andrew J Quinn, Benjamin AE Hunt, Matthew J Brookes, Anna C Nobre, and Mark W Woolrich. Spontaneous cortical activity transiently organises into frequency specific phase-coupling networks. Nature Communications, 9(1):2987, 2018.

[33] Gemeng Zhang, Biao Cai, Aiying Zhang, Julia M Stephen, Tony W Wilson, Vince D Calhoun, and Yu-Ping Wang. Estimating dynamic functional brain connectivity with a sparse hidden markov model. IEEE transactions on medical imaging, 39(2):488–498, 2019.

[34] Robyn L Miller, Maziar Yaesoubi, Jessica A Turner, Daniel Mathalon, Adrian Preda, Godfrey Pearlson, Tulay Adali, and Vince D Calhoun. Higher dimensional meta-state analysis reveals reduced resting fmri connectivity dynamism in schizophrenia patients. PLOS ONE, 11(3):e0149849, 2016.

[35] Ashkan Faghiri, Armin Iraji, Eswar Damaraju, Jessica Turner, and Vince D Calhoun. A unified approach for characterizing static/dynamic connectivity frequency profiles using filter banks. Network Neuroscience, 5(1):56–82, 2021.

[36] Zening Fu, Ishaan Batta, Lei Wu, Anees Abrol, Oktay Agcaoglu, Mustafa S Salman, Yuhui Du, Armin Iraji, Sarah Shultz, Jing Sui, et al. Searching reproducible brain features using NeuroMark: templates for different age populations and imaging modalities. Neuroimage, 292:120617, 2024.

[37] Y. Du, S. L. Fryer, D. Lin, J. Sui, J. Chen, E. Damaraju, E. Mennigen, B. Stuart, R. L. Loewy, D. H. Mathalon, J. M. Ford, S. McEwen, V. D. Calhoun, and S. Vinogradov. A novel spatially constrained ICA approach for robustly separating and identifying task-positive and task-negative networks from task fMRI data. Frontiers in Human Neuroscience, 10:599, 2016.

[38] Q. Lin, J. Liu, Y. Zheng, H. Liang, and V. D. Calhoun. Semiblind ICA of fMRI: A method for incorporating prior information into ICA algorithms. Journal of Neuroscience Methods, 192(1):98–112, 2010.

[39] Nora Leonardi, Jonas Richiardi, Markus Gschwind, Samanta Simioni, Jean-Marie Annoni, Myriam Schluep, Patrik Vuilleumier, and Dimitri Van De Ville. Principal components of functional connectivity: a new approach to study dynamic brain connectivity during rest. NeuroImage, 83:937–950, 2014.

[40] Anees Abrol, Eswar Damaraju, Robyn L Miller, Julia M Stephen, Eric D Claus, Andrew R Mayer, and Vince D Calhoun. Replicability of time-varying connectivity patterns in large resting state fMRI samples. NeuroImage, 163:160–176, 2017.

[41] K. L. Miller, F. Alfaro-Almagro, N. K. Bangerter, D. L. Thomas, E. Yacoub, J. Xu, A. J. Bartsch, S. Jbabdi, S. N. Sotiropoulos, J. L. R. Andersson, M. W. Woolrich, and S. M. Smith. Multimodal population imaging in the UK Biobank. Nature Neuroscience, 19(11):1523–1536, 2016.

[42] Thomas J Littlejohns, Jo Holliday, Lorna M Gibson, Steve Garratt, Niels Oesingmann, Fidel Alfaro-Almagro, Jimmy D Bell, Chris Boultwood, Rory Collins, Megan C Conroy, et al. The uk biobank imaging enhancement of 100,000 participants: rationale, data collection, management and future directions. Nature communications, 11(1):2624, 2020.

[43] Rongtao Jiang, Stephanie Noble, Matthew Rosenblatt, Wei Dai, Jean Ye, Shu Liu, Shile Qi, Vince D Calhoun, Jing Sui, and Dustin Scheinost. The brain structure, inflammatory, and genetic mechanisms mediate the association between physical frailty and depression. Nature Communications, 15(1):4411, 2024.

[44] David B Keator, Theo GM Van Erp, Jessica A Turner, Gary H Glover, Bryon A Mueller, Thomas T Liu, James T Voyvodic, Jerod Rasmussen, Vince D Calhoun, Hyo Jong Lee, et al. The function biomedical informatics research network data repository. Neuroimage, 124:1074–1079, 2016.

[45] Clifford R Jack, Matt A Bernstein, Nick C Fox, Paul Thompson, Gene Alexander, Danielle Harvey, Bret Borowski, Paula J Britson, Jennifer L Whitwell, Chadwick Ward, et al. The alzheimer’s disease neuroimaging initiative (ADNI): MRI methods. Journal of Magnetic Resonance Imaging, 27(4):685–691, 2008.

[46] Karl J. Friston, Andrew P. Holmes, Keith J. Worsley, Jean-Baptiste Poline, Christopher D. Frith, and Richard S. J. Frackowiak. Statistical parametric maps in functional imaging: A general linear approach. Human Brain Mapping, 2(4):189–210, 1995.

[47] Wellcome Centre for Human Neuroimaging. Statistical Parametric Mapping (SPM12). University College London, London, UK, 2014. Available at https://www.fil.ion.ucl.ac.uk/spm/software/spm12/.

[48] Kyle M Jensen, Jessica A Turner, Lucina Q Uddin, Vince D Calhoun, and Armin Iraji. Addressing inconsistency in functional neuroimaging: A replicable data-driven multi-scale functional atlas for canonical brain networks. biorxiv, 2024.

[49] M. Moein Esfahani, Vladislav Esaulov, Hemanth Venkateswara, and Vince Calhoun. Neuromark dfnc patterns: A fully automated pipeline to estimate subject-specific states from rs-fmri data via constrained ica of dfnc in +100k subjects. bioRxiv, 2025.

[50] Moein Esfahani, Hemanth Venkateswara, Zening Fu, Ram Ballem, and Vince Calhoun. Cnn and lstm models for fmri-based schizophrenia classification using c-ica of dfnc. medRxiv, 2025.

[51] Maria Giulia Preti, Thomas AW Bolton, and Dimitri Van De Ville. The dynamic functional connectome: State-of-the-art and perspectives. Neuroimage, 160:41–54, 2017.

[52] Karl J Friston. The disconnection hypothesis. Schizophrenia research, 30(2):115–125, 1998.

[53] William Pettersson-Yeo, Paul Allen, Stefania Benetti, Philip McGuire, and Andrea Mechelli. Dysconnectivity in schizophrenia: where are we now? Neuroscience & Biobehavioral Reviews, 35(5):1110–1124, 2011.

[54] Pinhong Chen, Enmao Ye, Xing Jin, Yiyu Zhu, and Ling Wang. Association between thalamocortical functional connectivity abnormalities and cognitive deficits in schizophrenia. Scientific Reports, 9:2952, 2019.

[55] Kyle M. Jensen, Tricia Z. King, Pablo Andrés-Camazón, Vince D. Calhoun, and Armin Iraji. Aberrant cortical–subcortical-cerebellar connectivity in resting-state fmri as an imaging marker of schizophrenia and psychosis: a systematic review of data-driven whole-brain functional connectivity analyses. Frontiers in Neuroimaging, Volume 4 – 2025, 2025.

[56] Kyle M Jensen, Vince D Calhoun, Zening Fu, Kun Yang, Andreia V Faria, Koko Ishizuka, Akira Sawa, Pablo Andrés-Camazón, Brian A Coffman, Dylan Seebold, et al. A whole-brain neuromark resting-state fmri analysis of first-episode and early psychosis: Evidence of aberrant cortical-subcortical-cerebellar functional circuitry. NeuroImage: Clinical, 41:103584, 2024.

[57] Neil D. Woodward, Haleh Karbasforoushan, and Stephan Heckers. Thalamocortical dysconnectivity in schizophrenia. American Journal of Psychiatry, 169(10):1092–1099, 2012.

[58] Monica Giraldo-Chica and Neil D. Woodward. Review of thalamocortical resting-state fmri studies in schizophrenia. Schizophrenia Research, 180:58–63, 2017.

[59] Maeri Yamamoto, Epifanio Bagarinao, Masanori Shimamoto, Tetsuya Iidaka, and Norio Ozaki. Involvement of cerebellar and subcortical connector hubs in schizophrenia. NeuroImage: Clinical, 35:103140, 2022.

[60] Simone Kühn and Jü rgen Gallinat. Quantitative meta-analysis on state and trait aspects of auditory verbal hallucinations in schizophrenia. Schizophrenia Bulletin, 38(4):779–786, 06 2012.

[61] Renaud Jardri, Alexandre Pouchet, Delphine Pins, and Pierre Thomas. Cortical activations during auditory verbal hallucinations in schizophrenia: a coordinate-based meta-analysis. American Journal of Psychiatry, 168(1):73–81, 2011.

[62] Daniel Collerton, James Barnes, Nico J. Diederich, Rob Dudley, Dominic ffytche, Karl Friston, Christopher G. Goetz, Jennifer G. Goldman, Renaud Jardri, Jaime Kulisevsky, Simon J.G. Lewis, Shigetoshi Nara, Claire O’Callaghan, Marco Onofrj, Javier Pagonabarraga, Thomas Parr, James M. Shine, Glenn Stebbins, John-Paul Taylor, Ichiro Tsuda, and Rimona S. Weil. Understanding visual hallucinations: A new synthesis. Neuroscience & Biobehavioral Reviews, 150:105208, 2023.

[63] Nancy C. Andreasen, Sergio Paradiso, and Daniel S. O’Leary. “cognitive dysmetria” as an integrative theory of schizophrenia: A dysfunction in cortical-subcortical-cerebellar circuitry? Schizophrenia Bulletin, 24(2):203–218, 01 1998.

[64] Yvette I. Sheline and Marcus E. Raichle. Resting state functional connectivity in preclinical alzheimer’s disease. Biological Psychiatry, 74(5):340–347, 2013.

[65] Katell Mevel, Gaël Chételat, Francis Eustache, and Béatrice Desgranges. The default mode network in healthy aging and alzheimer’s disease. International Journal of Alzheimer’s Disease, 2011:535816, 2011.

[66] Lisa T Eyler, Jeremy A Elman, Sean N Hatton, Sarah Gough, Anna K Mischel, Donald J Hagler, Carol E Franz, Anna Docherty, Christine Fennema-Notestine, Nathan Gillespie, et al. Resting state abnormalities of the default mode network in mild cognitive impairment: a systematic review and meta-analysis. Journal of Alzheimer’s Disease, 70(1):107–120, 2019.

[67] Sam Ereira, Sheena Waters, Adeel Razi, and Charles R Marshall. Early detection of dementia with default-mode network effective connectivity. Nature Mental Health, 2(7):787–800, 2024.

